# Local delivery of cell surface-targeted immunocytokines programs systemic anti-tumor immunity

**DOI:** 10.1101/2024.01.03.573641

**Authors:** Luciano Santollani, Yiming J. Zhang, Laura Maiorino, Joseph R. Palmeri, Jordan A. Stinson, Lauren R. Duhamel, Kashif Qureshi, Jack R. Suggs, Owen T. Porth, William Pinney, Riyam Al Msari, K. Dane Wittrup, Darrell J. Irvine

**Affiliations:** Department of Chemical Engineering, Massachusetts Institute of Technology; Cambridge, MA, USA; Koch Institute for Integrative Cancer Research, Massachusetts Institute of Technology, Cambridge; MA, USA; Department of Biological Engineering, Massachusetts Institute of Technology; Cambridge, MA, USA; Ragon Institute of Massachusetts General Hospital, Massachusetts Institute of Technology, and Harvard University; Cambridge, MA, USA; Howard Hughes Medical Institute; Chevy Chase, MD, USA; Department of Materials Science and Engineering; Massachusetts Institute of Technology, Cambridge, MA, USA

## Abstract

Cytokine therapies are potent immunotherapy agents but exhibit severe dose-limiting toxicities. One strategy to overcome this involves engineering cytokines for intratumoral retention following local delivery. Here, we develop a localized cytokine therapy that elicits profound anti-tumor immunity by engineered targeting to the ubiquitous leukocyte receptor CD45. We designed CD45-targeted immunocytokines (αCD45-Cyt) that, upon injection, decorated the surface of leukocytes in the tumor and tumor-draining lymph node (TDLN) without systemic exposure. αCD45-Cyt therapy eradicated both directly treated tumors and untreated distal lesions in multiple syngeneic mouse tumor models. Mechanistically, αCD45-Cyt triggered prolonged pSTAT signaling and reprogrammed tumor-specific CD8^+^ T cells in the TDLN to exhibit an anti-viral transcriptional signature. CD45 anchoring represents a broad platform for protein retention by host immune cells for use in immunotherapy.

## Main Text

Cytokines are a class of small proteins that serve as modulators of immune signaling cascades. Because of their multifunctional roles in lymphocyte migration, activation, and proliferation, cytokines have been long recognized as promising cancer immunotherapy agents(*1*–*3*). In fact, interleukin-2 (IL-2) was the first efficacious immunotherapy for advanced cancer, eliciting a 17% overall response rate as a monotherapy in metastatic melanoma leading to its FDA approval in 1998(*4*). Despite this clinical validation, dose-limiting toxicities and severe immune-related adverse events significantly hinder the clinical utilization of cytokines for oncology. This can be attributed to systemic lymphocyte activation triggered by the high doses that must be administered to achieve meaningful tumor concentrations.

Over the last 20 years, protein engineering has emerged as a tool to address the shortcomings of native cytokine therapies(*5*–*7*). One strategy to increase both safety and efficacy is to engineer cytokines for local delivery and retention within tumors and/or tumor-draining lymph nodes (TDLNs). We and others have previously demonstrated that potent cytokines and other immune agonists that exhibit high levels of toxicity when administered systemically can safely and effectively drive anti-tumor immunity when administered intratumorally (i.t.) to accessible lesions, by “anchoring” these drugs to intratumoral collagen, or to injected materials such as liposomes or alum particles(*8*–*15*). These approaches are enabled by modern interventional radiology methods that make intratumoral administration feasible in virtually any type of cancer(*15*–*17*). While these approaches elicit profound regressions of directly treated lesions, they typically elicit modest responses at distal untreated sites, often requiring an additional systemically administered agent such as checkpoint blockade immunotherapy (CBI) to drive abscopal effects. Limited abscopal responses are consistent with the findings of many other groups employing diverse therapeutic approaches in both preclinical models and early-stage clinical trials(*16*, *18*). Collectively, these data raise the question of whether localized immunotherapies are capable of eliciting robust systemic responses against metastatic disease.

Here we report a localized immunotherapy regimen targeting cytokines to the universal leukocyte receptor CD45. We apply this cell-surface targeting to two well-studied cytokines, IL-12 and IL-15. Optimizing dose level, tissue localization, and timing results in a highly efficacious and non-toxic cytokine therapy capable of eradicating treated tumors as well as eliciting complete responses at distal lesions in multiple syngeneic models. We characterize the response and mechanism of action of this therapy to show that CD45-targeted immunocytokines are retained on the cell-surface and trigger enhanced and prolonged signaling relative to their native cytokine counterparts. Leveraging this altered biology, we found that a single dose of CD45-targeted IL-12 followed by a single dose of CD45-targeted IL-15 dramatically altered T cell programming in the TDLN. This treatment expanded tumor-specific CD8^+^ T cells with potent effector phenotypes displaying a transcriptional signature mirroring effective responses to acute viral infection, while maintaining a pool of stem-like T cells. Dissemination of this potent effector T cell pool led to strong systemic anti-tumor immunity characterized by complete responses in both treated and untreated tumors in multiple syngeneic tumor models.

## RESULTS

### Engineering CD45 immunocytokines

We have previously shown that antibodies against the ubiquitous leukocyte surface receptor CD45 bind to the surface of T cells and other lymphocytes without triggering internalization(*19*, *20*). Analogous to recent reports of the altered biology achieved when IL-2 is targeted to lymphocytes via anti-PD-1(*21*), we hypothesized that cell surface-displayed cytokines using CD45 targeting might exhibit new biological effects relative to the native forms of these cytokines. To test this, we generated αCD45 immunocytokine fusions with IL-15 as a testbed payload. A murine superagonist IL-15 (IL-15 linked to a domain of its α receptor chain, IL-15Rα_sushi_) was fused to the C terminus of a murine IgG2c isotype heavy chain carrying LALA-PG effector attenuating mutations (hereafter, αCD45-IL15) (Fig. 1A)(*22*). We also synthesized a non-targeted size-matched control immunocytokine using an irrelevant fluorescein-specific antibody, referred to hereafter as IgG-IL15. We verified that recombinantly expressed αCD45-IL15 was able to bind plate-bound CD45 and trigger proliferation of IL-15 receptor-expressing reporter cells (Fig. 1, B and C, and fig. S1A).

**Fig. 1.**
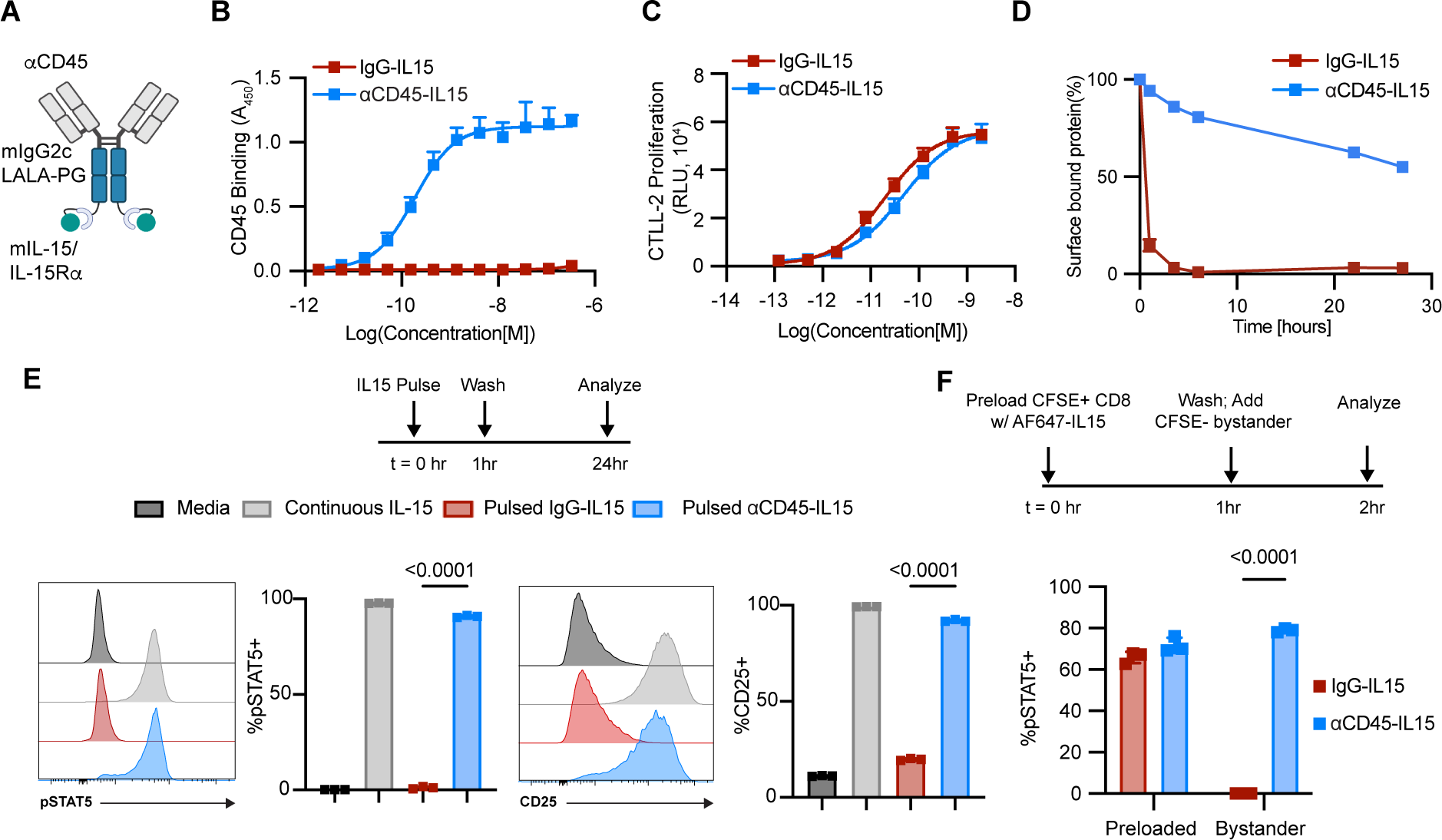
CD45 immunocytokines are retained on the cell surface and display extended signaling. **(A)** Schematic of IL-15 immunocytokine. **(B)** ELISA absorbance measurement of αCD45-IL15 and IgG-IL15 binding to plate-bound mouse CD45. **(C)** Luminescence measurement of CTLL-2 cell proliferation following 48 hr incubation with IgG-IL15 or αCD45-IL15 at indicated concentrations. **(D)** Internalization kinetics of IgG-IL15 or αCD45-IL15 labeled with AF488 following binding to primary activated CD8^+^ T cells. Surface signal was calculated by fluorescence quenching with an αAF488 antibody. **(E)** Primary CD8 T cells were pulsed for 1 hour with IL-15 immunocytokine fusions. Cells were washed to remove unbound cytokine, and 24 hours later fixed, permeabilized, and stained for flow cytometry analysis of pSTAT5 and CD25. **(F)** Primary CD8^+^ T cells were labeled with CFSE and incubated with AF647-labeled IL-15 immunocytokine fusions for 1 hr (“preloaded”), washed, and mixed into non-labeled “bystander” cells for 1 hr at a 1:20 preloaded: bystander ratio. Cells were then fixed, permeabilized, and stained for flow cytometry analysis of pSTAT5. Bystander cells were defined as CFSE^-^AF647^-^. All plots show mean ± SD from n = 3 biological replicates. P values were determined by one-way (E) or two-way (F) ANOVA followed by Tukey’s multiple comparison test.

Using dye-labeled constructs and fluorescence quenching, we assessed the binding and internalization behavior of these IL-15 immunocytokines incubated with activated primary CD8^+^ T cells. Control IgG-IL15 fusions bound to the cells and were rapidly internalized, while αCD45-IL15 had a cell-surface half-life of 24 hours, suggesting that CD45 binding can maintain cytokines on the cell surface irrespective of the rapid internalization typical for the native cytokine (Fig. 1D, and fig. S1B). CD45-targeted IL-15 bound to T cells at far higher levels than IgG-IL15, suggesting that CD45 increases the immunocytokine target receptor pool due to its high expression level (fig. S1C). To evaluate whether this extended surface retention altered cytokine activity, we pulsed primary CD8^+^ T cells with IL-15 immunocytokines for 1 hr, then washed to remove unbound cytokine and evaluated downstream signaling over time. Strikingly, T cells briefly pulsed with αCD45-IL15 exhibited robust pSTAT5 and CD25 expression levels 24 hours later that were almost as high as cells incubated continuously in IL-15 (Fig. 1E). By contrast, these markers of IL-15 signaling had all returned to near baseline by this timepoint for cells pulsed with non-targeted IgG-IL15 (Fig. 1E). We next tested whether CD45-anchored cytokines could signal to neighboring cells *in trans*. CFSE-labeled CD8^+^ T cells were pre-loaded with AlexaFluor-labeled αCD45-IL15 by pulsing for 1 hour with the construct, then washed and mixed with non-loaded CFSE^-^ “bystander” CD8^+^ T cells for an additional hour. The presence of just 1 pre-loaded T cell per 20 total cells led to pSTAT5 induction in a majority of both the preloaded and bystander cultures, though at a higher level in the former (Fig. 1F and fig. S1D). Signaling to bystander cells occurred with minimal detectable transfer of the labeled cytokine fusion to the bystander cells, suggesting prominent *in trans* signaling of the cytokine from pre-loaded to bystander T cells (fig. S1E). By contrast, non-targeted IgG-IL15 led to robust pSTAT5 signal in the preloaded population but failed to stimulate any bystander cells, likely due to its rapid internalization (Fig. 1F). To probe whether CD45 surface retention is a mouse-specific phenomenon, we performed internalization assays with human CD8^+^ T cells and an anti-human CD45 antibody, and found, similar to the murine antibody, prolonged cell surface retention with a half of life of over 5 days (fig. S1F). To determine if cell surface retention and prolonged signaling was a general phenomenon for αCD45-cytokine fusions, we also generated αCD45-IL12, a fusion of a single chain IL12p70 with αCD45. This IL-12 immunocytokine was able to bind CD45 with no loss of cytokine bioactivity and led to increased surface retention as well as extended signaling, as measured by prolonged pSTAT4 levels (fig. S2, A to E). Thus, although CD45’s phosphatase properties have been previously employed to deactivate nearby tethered receptors(*23*), CD45-targeted antibody-cytokine fusions as designed here exhibit sustained and potent signaling. Together, this suggests that endowing cytokines with the ability to bind CD45 in addition to their cognate receptor modulates downstream signaling by slowing internalization.

### Local αCD45-IL15 persists at the tumor and TDLN

We next assessed the *in vivo* biodistribution of these fusion proteins following intratumoral administration, reasoning that binding to leukocytes in the tumor and TDLN could allow for strong local immune stimulation without the toxicity observed with systemic cytokine administration. AlexaFluor-labeled IL-15 immunocytokines were injected into established MC38 flank tumors, and association of the constructs on cells in the tumor, TDLNs, and distribution into the blood 24 hours later was assessed by flow cytometry, microscopy, and fluorescence measurements (Fig. 2A). Injection of 10 µg αCD45-IL15 led to robust labeling of a majority of all CD45^+^ cells in the tumor (Fig. 2, B to D). By contrast, the non-targeted control cytokine only showed association with ∼40% of intratumoral CD45^+^ cells, and at much lower MFI levels than the αCD45 fusion (Fig. 2, B to D). αCD45-IL15 bound effectively to all common intratumoral immune subsets while IgG-IL15 mostly associated with Natural Killer (NK) cells as the dominant target, likely due to high IL2RβƔ expression by NK cells (fig. S3A and fig. S4). Confocal microscopy confirmed these results, showing broad distribution of αCD45-IL15 throughout the tumor (Fig. 2E). IgG-IL15 exposure was sparse and isolated to narrow sections of the tumor (Fig. 2E). This dose also led to labeling of ∼75% of CD45^+^ cells in the TDLN by αCD45-IL15, whereas IgG-IL15 labeling was not statistically significant relative to untreated controls (Fig. 2F and fig. S3B). Microscopy imaging revealed efficient drainage of labeled αCD45-IL15 throughout the TDLN and was readily apparent on CD8^+^ T cells (Fig. 2G and fig. S3C). Profiling of cytokine uptake by immune cell subsets in the TDLN again revealed NK cells as the only cellular subset with substantial uptake of IgG-IL15, while αCD45-IL15 was bound to T cells, B cells, macrophages and dendritic cells (fig. S3D). Importantly, at this dose, no free αCD45-IL15 was detected in the serum and there was minimal association with peripheral blood mononuclear cells (PBMCs), while non-targeted IgG-IL15 was found accumulating in the blood (Fig. 2H and fig. S3E). Intratumoral administration was required for simultaneous tumor and lymph node labeling, as peritumorally delivered αCD45-IL15 efficiently labeled the TDLN but failed to meaningfully accumulate in the tumor (fig. S3F). Thus, CD45 targeting increased localization of the cytokine on immune cells in both tumors and TDLNs, while avoiding systemic exposure when doses were selected to not exceed the binding capacity of leukocyte surface CD45 in the tumor and TDLN.

**Fig. 2.**
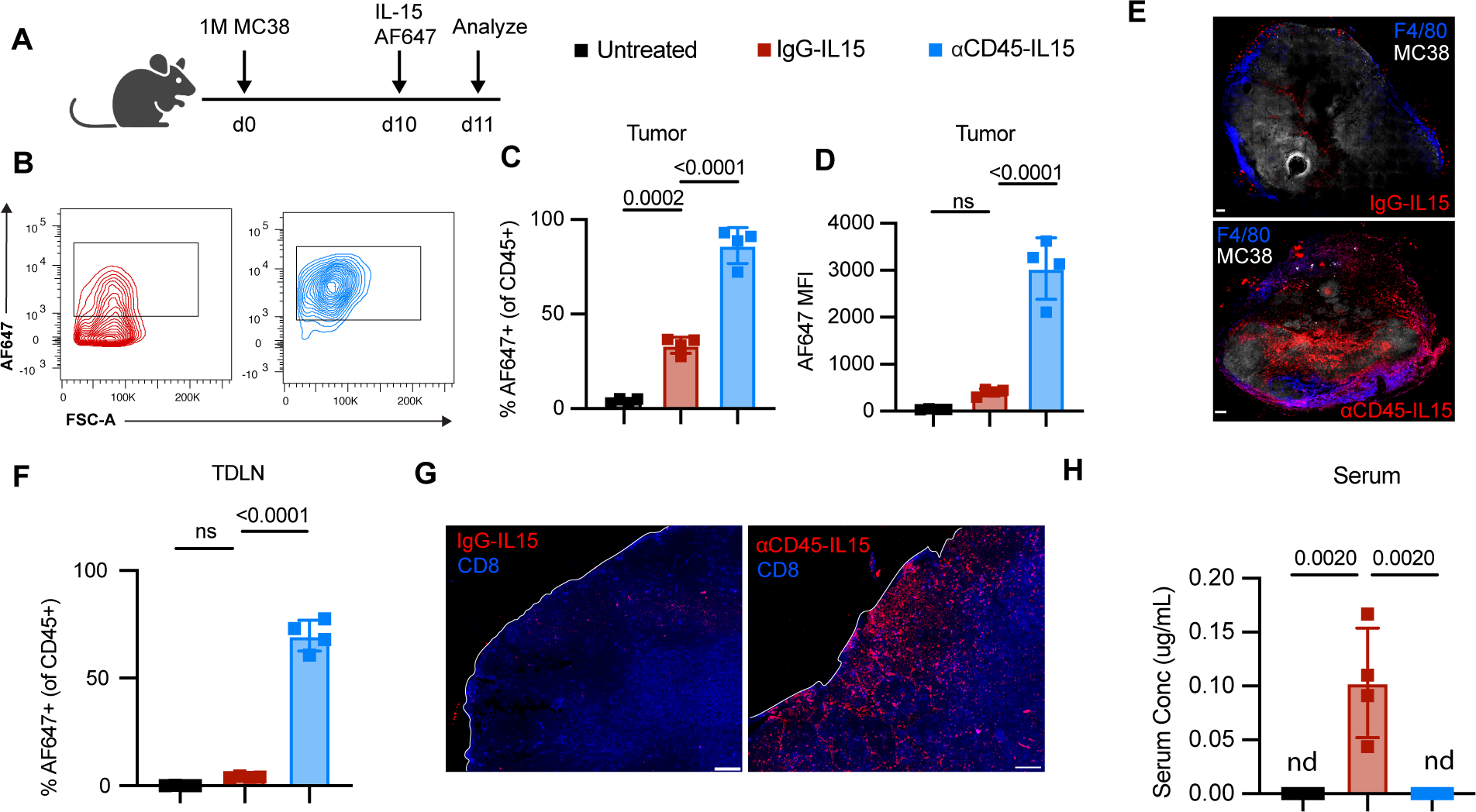
Intratumorally administered αCD45-IL-15 is retained in the tumor and TDLN with negligible systemic exposure. C57BL/6J mice (*n* = 4 mice/group) were inoculated with 1M MC38 or MC38-ZsGreen cells in the flank, and 10 days later administered AF647-labeled αCD45-IL15 or IgG-IL15 (10µg) intratumorally, followed by analysis of cytokine biodistribution 24 hr later. For microscopy experiments, MC38-ZsGreen tumors were used for easy detection of tumor cells. **(A)** Study diagram. **(B)** Representative histograms of antibody-cytokine fusion binding to CD45^+^ tumor-infiltrating leukocytes. **(C)** Frequencies of AF647^+^ tumor-infiltrating leukocytes. **(D)** AF647 MFI on TILs. **(E)** Representative confocal images of tumors. Scale bar, 200μm . **(F)** AF647 labeling on TDLN immune cells. **(G)** Representative histological images of TDLN. Scale bar, 100 μm. **(H)** Serum concentrations of dosed immunocytokines based on fluorescence spectroscopy measurements. nd = not detected. All plots show mean ± SD from *n* = 4 replicates/group. P values were determined by one-way ANOVA followed by Tukey’s multiple comparison test.

### αCD45-cytokine therapy eradicates treated tumors

Next, we evaluated the impact of this altered signaling and biodistribution on the therapeutic potential of a cytokine therapy employing both αCD45-IL15 and αCD45-IL12. We first treated established tumors 30-35 mm^2^ in size with a single i.t. injection of 1 µg αCD45-IL12 followed by one injection of 10 µg αCD45-IL15 several days later. Our rationale for this sequencing was to use the IL-12 immunocytokine to inflame the tumor and promote an initial wave of tumor immunogenic cell death and tumor antigen release by pre-existing or newly recruited TILs(*12*), followed by amplification of newly primed tumor-specific T cells in the TDLN by the IL-15 immunocytokine several days later(*24*, *25*). The selected doses were informed by findings from the biodistribution experiments and aimed to retain IL-12 at the tumor but deliver IL-15 to both the tumor and TDLN. On a cytokine mass basis, these doses did not surpass 2 µg of total cytokine. Hereafter, we refer to this sequential αCD45-IL12/αCD45-IL15 regimen as αCD45-Cyt; the untargeted cytokine control therapy consisting of IgG-IL12 followed by IgG-IL15 is referred to as IgG-Cyt.

We tested this therapeutic regimen in two common syngeneic tumor models, the MC38 colorectal carcinoma, which is partially responsive to checkpoint blockade immunotherapy (CBI), and B16F10, an aggressive and poorly responsive melanoma model (Fig. 3A). Injection of control IgG-Cyt elicited an extension in survival, but few (MC38) or no (B16F10) animals were long-term survivors (Fig. 3A and fig. S5, A and B). By contrast, 100% (MC38) or >90% (B16F10) of tumors treated with αCD45-Cyt therapy were rejected, and animals exhibited no signs of cytokine toxicity, as assessed via weight loss (Fig. 3A and fig. S5, A and B). Potential modulation of the immune response by CD45 engagement did not contribute to efficacy, as treatment with αCD45 in the absence of cytokines provided no tumor control (fig. S5C). We assessed the mechanism of tumor rejection by αCD45-Cyt therapy through antibody depletions and knockout mice: αCD45-Cyt therapy entirely depended on CD8^+^ T cells and *batf3^+^* dendritic cells, consistent with a critical role for tumor antigen cross-presentation (Fig. 3B). NK cells, neutrophils, and macrophages were each dispensable for anti-tumor efficacy (Fig. 3B). Mice cured of MC38 and B16F10 tumors by αCD45-Cyt therapy displayed robust immune memory upon secondary tumor rechallenge (Fig. 3C and fig. S5D). Use of both cytokines in the treatment was required, as αCD45-IL15 or αCD45-IL12 monotherapy using the same treatment schedule elicited no long-term survivors (fig. S5E).

**Fig. 3.**
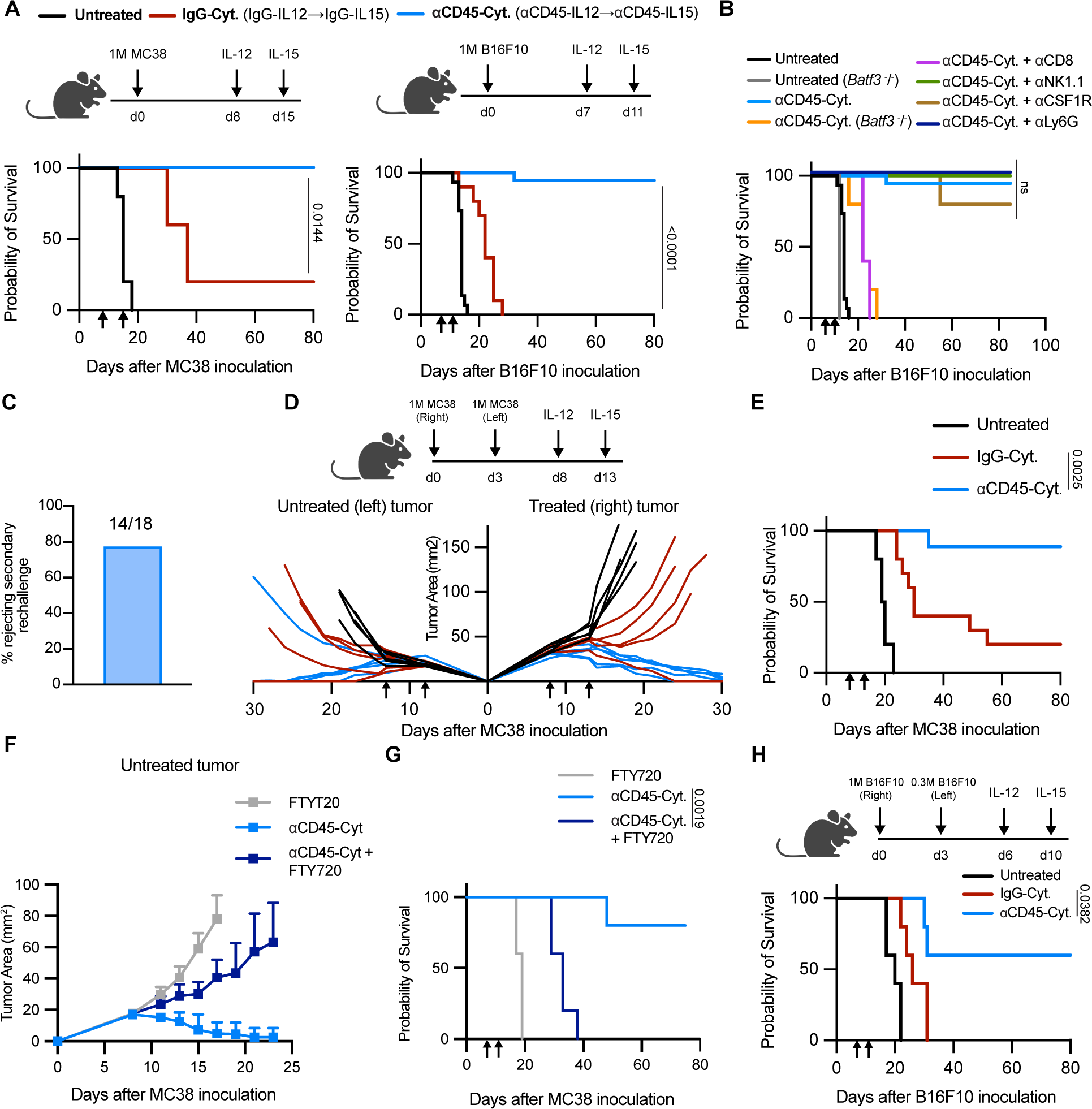
αCD45-cytokine therapy eradicates injected tumors and primes robust systemic anti-tumor responses. **(A)** Kaplan Meier survival of mice (*n* = 5/group for MC38, n = 10-19/group for B16F10 across 2-5 independent experiments) inoculated with 1M tumor cells and treated with 1 µg of αCD45-IL12 and 10µg of αCD45-IL15 (referred to as αCD45-Cyt) or treated with 1 µg of IgG-IL12 and 10µg of IgG-IL15 (referred to as IgG-Cyt) as shown on the experimental timelines. Statistical comparison shown between IgG-Cyt and αCD45-Cyt groups. **(B)** Kaplan Meier survival of mice (*n* = 5/group) bearing B16F10 tumors treated with αCD45-Cyt therapy in combination with depleting antibodies or performed in knockout mice as shown. **(C)** Overall survival of mice cured of B16F10 tumors rechallenged with 0.1M B16F10 cells 100 days after the initial challenge. **(D)** Mice bearing MC38 tumors on both flanks were treated with 1µg IL-12 and 10µg of IL-15 in the right tumor only at indicated times (*n* = 9-10 mice/group across 2 independent experiments). Shown are individual tumor growth curves from one representative experiment. **(E)** Kaplan Meier survival from experiment shown in (D). **(F)** Mice bearing bilateral MC38 tumors were treated with αCD45-Cyt therapy in the presence of FTY720 starting on d5. Shown is the average tumor growth curve for the untreated tumor. **(G)** Kaplan Meier survival from experiment shown in (F). **(H)** Kaplan Meier survival of mice bearing contralateral B16F10 tumors treated with 1 µg IL-12 and 10µg of IL-15 as shown (*n*=5 mice/group). P values were determined by log-rank (Mantel-Cox) test. For all plots, arrows indicate treatment.

To study the importance of cytokine localization to the tumor and TDLN, we varied the route of administration of αCD45-Cyt therapy. Dosing the same αCD45-Cyt treatment paradigm systemically (i.p.) led to a complete loss of efficacy, confirming the importance of intratumoral exposure and efficient TDLN drainage for activity (fig. S5, F to H). Next, to determine the relevant contributions to efficacy from tumor vs TDLN leukocytes, we administered αCD45-Cyt therapy peritumorally, which predominantly acts on the TDLN while sparing the tumor (fig. S3E). While dosing both cytokines peritumorally completely ablated efficacy, we found that intratumoral αCD45-IL12 followed by peritumorally delivered αCD45-IL15 displayed comparable early tumor control to the full intratumoral paradigm (fig. S5I). However, a majority of these mice (3/5) eventually succumbed to tumor burden, suggesting that optimal long-term tumor control and rejection required both doses to be administered intratumorally (fig. S5J).

### αCD45-cytokine therapy elicits systemic immunity

The successful translation of intratumorally administered immunotherapies will depend on their ability to drive a systemic anti-tumor immune response that can eliminate lesions that are not directly treated. To evaluate abscopal immune responses elicited by this localized therapy, MC38 tumors were inoculated on opposite flanks of mice followed by treatment of only the right-flank tumor with αCD45-Cyt therapy. Strikingly, this led to 90% of mice rejecting both the treated and untreated tumors and becoming long-term survivors, while only 20% of animals receiving IgG-Cyt therapy eliminated both tumors (Fig. 3, D and E). These results were obtained despite the fact that tracking of fluorescently-labeled constructs showed that locally delivered IgG-Cyt readily diffused out of the treated lesion and accumulated in the distal site, while no statistically significant accumulation of αCD45-Cyt was measured at the untreated tumor (fig. S6A). To determine whether regression of distal tumors relied on lymphocyte trafficking, we tested αCD45-Cyt in the same two-tumor MC38 model but in the presence of FTY720 to block lymphocyte exit from lymphoid tissue (fig. S6B). The addition of FTY720 ablated the ability of αCD45-Cyt to control the untreated left-flank tumor and led to no long-term survivors, suggesting that rejection of untreated lesions is driven by lymphocyte migration (Fig. 3, F and G). αCD45-Cyt therapy still triggered regression of the treated tumor in the presence of FTY720, but no tumors had been cured at the time of euthanasia due to outgrowth of the untreated distal lesion (fig. S6C). To further stress test this treatment, we tested abscopal responses in the aggressive B16F10 model. While IgG-Cyt treatment led to some delay of treated tumors, all untreated tumors progressed (Fig. 3H and fig. S6D). By contrast αCD45-Cyt therapy led to primary tumor regression in all mice and only 2 of 5 untreated tumors escaped (Fig. 3H and fig. S6D). Finally, to assess the ability of αCD45-Cyt therapy to eliminate metastases, we treated mice that had been injected with 0.2M B16F10 cells intravenously in addition to flank tumor inoculation, giving rise to lung metastases. Once untreated mice had reached euthanasia criteria due to flank tumor outgrowth (day 15), we analyzed lung metastasis control across groups. In line with our previous experiments, regression of the treated primary tumor was in progress for αCD45-Cyt therapy at this time point. We also found significant control of lung metastases by αCD45-Cyt therapy, as measured by a dramatic reduction in total metastatic tumor burden and average metastasis size (fig. S7, A to D). Altogether, we find that αCD45-Cyt therapy is able to prime a robust systemic immune response after localized treatment that is able to control both treated and distal untreated lesions.

### Treated TDLN CD8^+^ T cells exhibit transcriptional signatures mirroring an acute anti-viral response

We next sought to understand how localized αCD45-Cyt treatment elicited such potent systemic immunity, focusing on the CD8^+^ T cells required for efficacy. Recent reports demonstrating the spatial segregation of T cell priming and activation between the TDLN and tumor(*26*), respectively, motivated us to profile the TDLN CD8^+^ phenotypes after treatment. To assess effects of αCD45-Cyt therapy on tumor-specific T cells, we treated B16F10 tumors with αCD45-Cyt or IgG-Cyt therapy, and 1 day after dosing of the IL-15 immunocytokine, sorted CD8^+^ T cells specific for the immunodominant p15E endogenous retroviral antigen expressed by B16F10 tumors(*27*) via peptide-MHC tetramer staining for downstream RNA sequencing (RNA-seq). αCD45-Cyt therapy dramatically reprogrammed the CD8^+^ T cell response, leading to significant upregulation of 1726 genes and downregulation of 1279 genes in comparison to the untreated condition (Fig. 4A). Many upregulated genes were associated with effector function (*IL2ra, Gzmb, Klrg1, Prf1*) as well as interferon signaling (*Ifng*, *Ifngr1*, *Ifitm1*), suggesting an ability of αCD45-Cyt therapy to promote activation and effector differentiation of T cells within the TDLN (Fig. 4A). Concurrently, genes related to lymph node trafficking (*Cxcr3, Cxcr5*) as well as stemness (*Tcf7*) were downregulated (Fig. 4A).

**Fig. 4.**
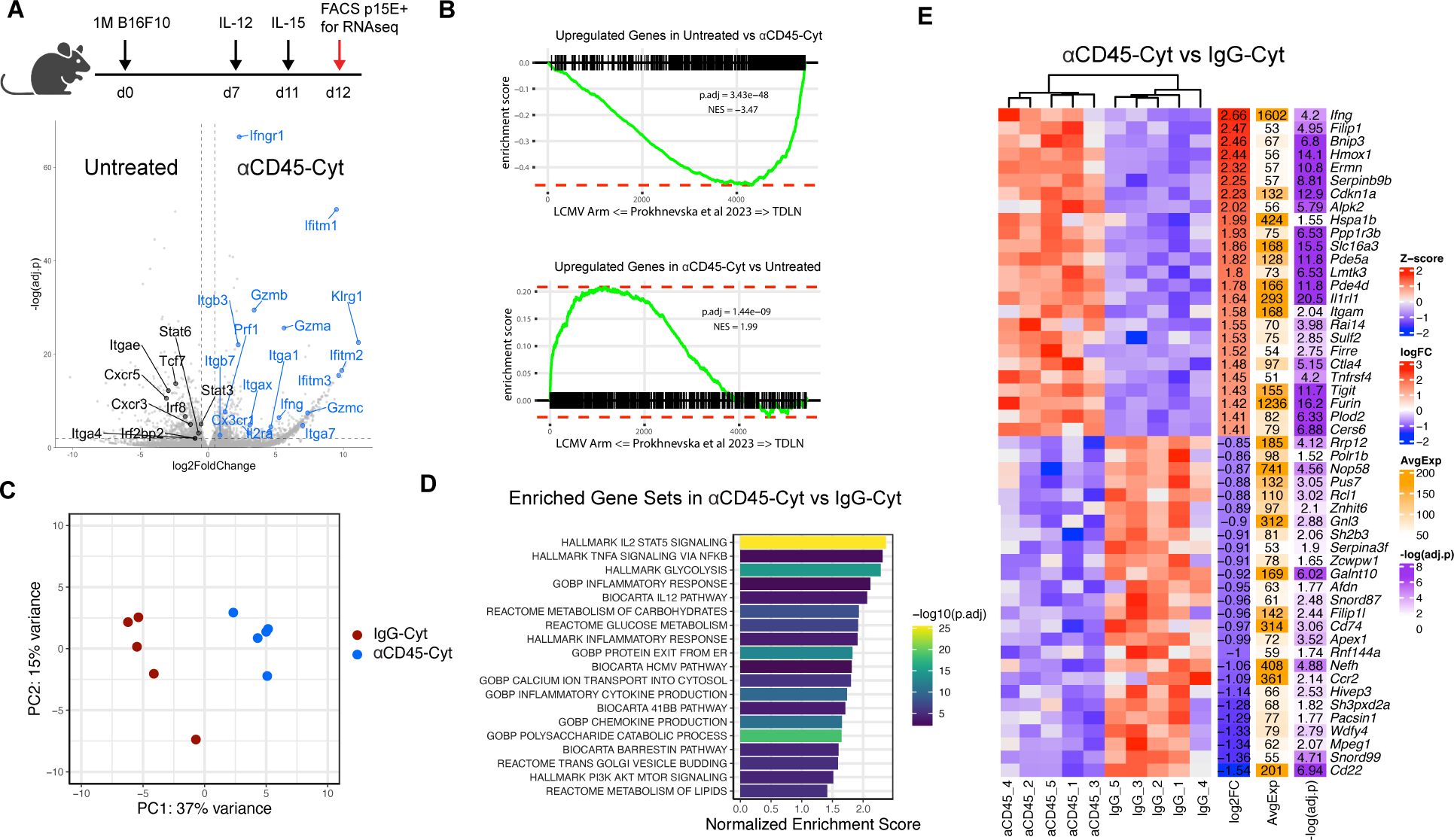
αCD45-Cyt therapy induces a viral-like signature in the tumor-specific TDLN compartment. B16F10 tumors (n = 4 or 5/ group) treated with αCD45-Cyt or IgG-Cyt therapy were sorted for CD8^+^ T cells specific for the immunodominant p15E retroviral antigen for downstream RNA-seq. **(A)** Volcano plot of differentially expressed genes between untreated and αCD45-Cyt with selected genes labeled. **(B)** GSEA using the differentially expressed genes between untreated and αCD45-Cyt, compared with T cell gene signatures from LCMV-Arm infection or TDLN from Prokhnevska et al. 2023 (GSE216731). Enrichment score is plotted. Adjusted p values and normalized enrichment scores (NES) are labeled. **(C)** PCA of IgG-Cyt and αCD45-Cyt treated samples. **(D)** Enriched pathways in αCD45-Cyt samples compared to IgG-Cyt samples. **(E)** Top 25 differentially expressed genes between αCD45-Cyt samples and IgG-Cyt samples. Statistical significance was determined using the Wald test with Benjamini-Hochberg post hoc correction.

Recent reports have demonstrated that, unlike the endogenous anti-tumor response, effective virus-specific T cell responses, such as following acute lymphocytic choriomeningitis virus Armstrong (LCMV Arm) infection, are accompanied by high expression of inflammatory cytokines and differentiation markers in responding lymph nodes(*26*). We thus performed gene set enrichment analysis (GSEA) to probe whether αCD45-Cyt therapy induces a “viral-like” signature within the tumor-reactive TDLN CD8^+^ compartment. We compared the untreated and αCD45-Cyt treated transcriptomes of tumor-specific CD8^+^ T cells for enrichment of a previously reported tumor (“TDLN”) or acute viral (“LCMV Arm”) transcriptional response (Fig. 4B). Transcriptional profiles of CD8^+^ T cells from untreated TDLNs closely matched the previously reported endogenous anti-tumor response, broadly characterized by a lack of effector molecules and activation markers (Fig 4B and fig. S8A). In contrast, αCD45-Cyt therapy elicited a signature highly enriched for the canonical anti-viral response, suggesting that the prolonged cytokine exposure within TDLNs induced by αCD45-Cyt triggers an immune response mirroring effective anti-viral T cell priming (Fig. 4B and fig. S8A). We next characterized differences in the transcriptional response of αCD45-Cyt treatment vs. control IgG-Cyt therapy. Principal-component analysis (PCA) showed that IgG-Cyt and αCD45-Cyt transcriptomes clustered separately (Fig. 4C). Pathway analysis comparing the two treatment conditions revealed a striking upregulation of hallmark STAT5 signaling triggered by αCD45-Cyt therapy, likely driven by the extended signaling of CD45-bound IL-15, as shown in Fig. 1E (Fig. 4D). Other upregulated genes belonged to pathways involving inflammatory responses, cytokine signaling, and cell-cycle metabolism (Fig. 4D). Finally, we examined the top 20 differentially expressed genes (DEGs) between αCD45-Cyt and IgG-Cyt therapy and found *Ifng* expression as the most differentially upregulated, along with other metabolism and inflammation genes (Fig. 4E and fig. S8B). Altogether, these data are indicative of a highly activated and proliferative TDLN compartment following αCD45-Cyt therapy.

### αCD45 immunocytokines generate optimal effectors

Recently, genetically-induced sustained STAT5 activation has been shown to rewire exhausted T cells into a unique effector profile that enables control of chronic infection or tumors(*28*). Based on our pathway analysis and *in vitro* data, we hypothesized that αCD45-IL15 stimulation may therapeutically elicit similar sustained STAT5 induction and downstream generation of tumor-specific effectors, which could then traffic to distal lesions and promote tumor rejection. Probing the signaling dynamics of CD45 immunocytokines in the TDLN 24 hours after intratumoral dosing with αCD45-IL15 alone revealed robust STAT5 phosphorylation induced in 80% of CD8^+^ T cells in the TDLN (Fig. 5, A and B). By contrast, injection of IgG-IL15 led to a dim pSTAT5^+^ population, closely mirroring the *in vitro* and RNA-seq results shown in Figs. 1E and 4D (Fig. 5, A and B). Next, using an identical experimental set-up to that used in the RNA-seq analysis, we treated B16F10 tumors with αCD45-Cyt or IgG-Cyt therapy and profiled p15E^+^CD8^+^ T cells in the tumor and TDLN 24 hours after IL15 dosing by flow cytometry. Only αCD45-Cyt treatment led to a statistically significant expansion of tetramer positive CD8^+^ T cells in the TDLN relative to untreated tumors (Fig. 5, D and E). In agreement with the RNA-seq results, in addition to expanded numbers, TDLN tumor-specific cells primed by αCD45-Cyt exhibited a profound increase in IFNγ expression as measured by MFI and the number of IFNγ^+^p15E^+^ cells (Fig. 5, F and G, and fig. S9A). IgG-Cyt therapy was unable to elicit IFNγ production within the TDLN (Fig. 5, F and G, and fig. S9A). Tumor-reactive cells in the TDLN additionally displayed high levels of Granzyme B expression after treatment with either cytokine therapy (fig. S9B). Both cytokine therapies also significantly increased CD25 expression over untreated control, but αCD45-Cyt treatment induced this upregulation on a larger fraction of the CD8^+^ compartment (fig. S9C).

**Fig. 5.**
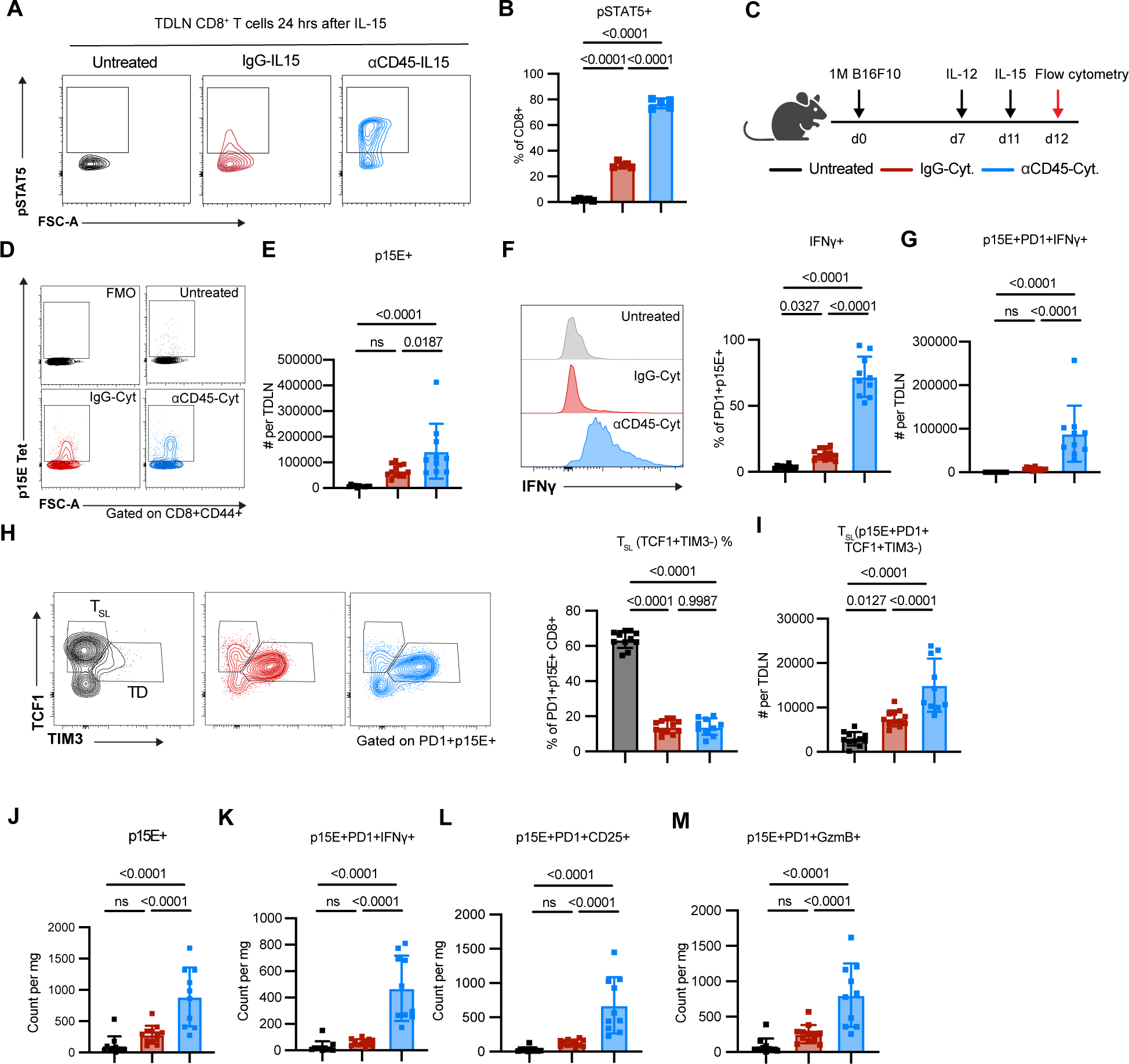
αCD45-cytokine therapy reprograms the TDLN leading to an optimal tumor-specific CD8^+^ T cell effector subset capable of infiltrating the TME. (**A** and **B**) C57BL/6J mice (*n*=5/group) inoculated with B16F10 tumors were treated on d7 with 10 μg IL-15 immunocytokine. 24 hours later, lymph nodes were fixed, permeabilized, and stained for pSTAT5. (A) Representative pSTAT5 contour plots after IL-15 treatment, previously gated on live CD3^+^CD8^+^ cells. (B) Summary data for pSTAT5 induction shown in (A). (**C** to **M**) Mice (*n* = 10-12/group across two independent studies) inoculated with 1M B16F10 tumor cells were treated with IgG-Cyt or αCD45-Cyt therapy starting on d7. Lymph nodes and tumors were harvested 24 hours after completion of therapy. (D) Representative contour plots and gating for p15E tetramer^+^ cells in the TDLN, previously gated on live CD8^+^CD44^+^ cells. (E) Treatment effects on p15E^+^ tumor-reactive cells in TDLN. (F) (Left) sample histogram of IFNγ expression and (right) TDLN frequency of tumor-specific cells expressing IFNγ^+^. (G) TDLN counts of IFNγ^+^PD-1^+^p15E^+^ T cells. (H) (Left) stem-like (T_SL_) vs. terminally differentiated (TD) sample gating and (right) TDLN frequency of stem-like PD1^+^TCF1^+^CD8^+^ T cells. (I) TDLN counts of stem-like PD1^+^TCF1^+^CD8^+^ T cells. (J) Counts of p15E tetramer^+^CD8^+^ TILs. (K) Counts of p15E tetramer^+^CD8^+^ TILs expressing IFNγ. (L) Counts of p15E tetramer^+^CD8^+^ TILs expressing CD25. (M) Counts of p15E tetramer^+^CD8^+^ TILs expressing Granzyme B. All plots show mean ± SD. P values were determined by one-way ANOVA followed by Tukey’s multiple comparison test.

Recent studies have suggested that antigen-specific TCF1^+^ stem-like T cells in disease-site-draining lymph nodes (in models of tumors, chronic infections, or autoimmune disease) play a critical role as a source of effector cell generation(*26*, *28*–*33*). Given the robust effector response exhibited in treated TDLNs, we evaluated the impact of αCD45-Cyt therapy on the magnitude of the stem-like (T_SL_, TCF1^+^TIM3^-^) compartment. Though both cytokine therapies led to the expected drop in the relative proportion of TCF1^+^ T cells, the absolute number of tumor-specific stem-like cells expressing PD1^+^ and TCF1^+^ increased, with αCD45-Cyt therapy eliciting a 6-fold higher count of stem-like cells in the TDLN over untreated mice (Fig. 5, H and I). Conversely, cytokine therapy skewed CD8^+^ T cells towards a terminally differentiated (TD, TCF1^-^TIM3^+^) phenotype as seen by high TIM3 expression in both treatment groups (fig. S9D). However, αCD45-Cyt therapy triggered a significantly larger expansion of the TD compartment by absolute count over IgG-Cyt therapy (fig. S9E).

Analysis of intratumoral p15E^+^CD8^+^ T cells revealed similar patterns as in the TDLN. αCD45-Cyt therapy led to increased numbers of tumor-reactive CD8^+^ T cells not seen after IgG-Cyt therapy (Fig. 5J). Phenotypic profiling once again highlighted the unique ability of αCD45-Cyt, but not IgG-Cyt, therapy to induce significantly higher levels of IFNγ within p15E-specific cells (fig. S9F). This led to significantly higher numbers of intratumoral IFNγ^+^CD8^+^cells (Fig. 5K). Mice treated with αCD45-Cyt also displayed higher levels of CD25 expression, both by percentage and magnitude (Fig. 5L and fig. S9G). Finally, intratumoral Granzyme B expression was high across all groups, but only αCD45-Cyt therapy was able to generate a significant increase in the compartment size (Fig. 5M and fig. S9H). Taken together, these immunophenotyping results show that local αCD45-Cyt therapy is not only able to dramatically alter the treated TME, as other local therapies have accomplished, but also reprogram the tumor-specific CD8^+^ population in the TDLN, correlating with stronger abscopal responses.

## Discussion

Antibody-targeted cytokines have been pursued for many years with modest impact, driven in part by the focus on targeting cancer cells lacking truly disease-specific antigens and the problem that their biodistribution is often governed by the cognate cytokine receptor and not the antibody target(*5*, *34*). Recent approaches, focusing on targeting immune cells through lineage markers or phenotype have shown promising preclinical data(*21*, *35*). A report of a PD-1 targeted IL-2 immunocytokine demonstrated unique internalization and signaling behavior relative to the parental cytokine, highlighting the potential of multi-receptor targeting to elicit novel cell signaling biology(*21*). Here we focused on targeting host immune cells, using the unique biology of CD45 to develop αCD45-cytokine fusions capable of modulating tumor-specific T cells within the TME and TDLN. We leveraged CD45 as a unique cell-based “anchor” for cytokines in two key ways: 1) its ability to be bound without triggering internalization, and 2) its abundant expression on the immune cell surface, with reported measurements upwards of 100,000 molecules per cell(*36*). We find evidence that these two characteristics allow for both *cis* and *trans* signaling of CD45-tethered cytokines, as shown by the ability of CD8 T cells preloaded with αCD45-IL15 to efficiently signal to multiple unloaded, bystander cells. Thus, by decorating leukocytes *in vivo* with CD45-targeted immunocytokines, bystander cells lacking the cognate receptor can serve as cellular depots that “present” the cytokine to nearby cells, extending the cytokine’s residence time. We show that specified intratumoral doses of CD45-targeted IL-15 and IL-12 are retained at the tumor and TDLN for prolonged periods of time, leading to extended pSTAT5 signaling and acquisition of a potent effector program by tumor-reactive CD8^+^ T cells. When we applied this strategy in a therapeutic context, this led to a reprogramming of the anti-tumor CD8^+^ T cell response at the TDLN and robust systemic anti-tumor immunity. On a cytokine basis, the doses given in the αCD45-Cyt paradigm did not exceed 2 µg, a highly dose-sparing regimen highlighting the benefit of administration at the right time and place.

Recent studies have suggested the endogenous cancer response is a two-step priming and activation process that is confined to TDLN and tumors, respectively(*26*). However, treatment with αCD45-Cyt therapy was able to alter this endogenous response and triggered a striking expansion of activated cytotoxic effector cells within the TDLN characterized by high expression of CD25, IFN-γ, granzyme B, and TIM3. In striking contrast to the endogenous unmanipulated anti-tumor response, T cells within the αCD45-Cyt-treated TDLN had transcriptional signature resembling an acute viral response. These changes in T cell activation in the TDLN also have many similarities with effects recently reported for antigen-specific T cells transduced to express constitutively active STAT5 (STAT5CA) in the setting of chronic LCMV infection(*28*). In that case, splenic STAT5CA T cells were found to transition to a polyfunctional effector state, with depletion of the exhausted precursor TCF1^+^ subset, and persist in the face of chronic infection. Interestingly, prolonged signaling from cell-bound αCD45-IL15 may represent a pharmacologically feasible approach to achieve similar effects in native T cells. Importantly however, αCD45-Cyt therapy does not appear to deplete the tumor-specific stem-like TCF1^+^PD1^+^ subset that is thought to be the critical self-renewing precursor to produce effector cells.

The ability to generate these potent effectors at the TDLN, in addition to the tumor, may explain the systemic response primed by αCD45-Cyt therapy. We hypothesize that tumor-reactive CD8^+^ T cells in TDLNs acted on by αCD45-Cyt therapy migrate into the systemic circulation and disseminate agnostically to distal lesions. In our treatment paradigm, we found evidence for a robust systemic response in two-tumor models of both MC38 colon carcinoma and B16F10 melanoma, as well as in a metastasis model using B16F10 lung metastases in combination with a flank tumor. These abscopal responses were not driven by drug leakage into the contralateral tumor, but were dependent on immune cell trafficking, as FTY720 ablated tumor control at the untreated lesion. These results suggest that nodal cytokine retention following administration can prime potent anti-tumor immunity capable of systemic tumor control.

This approach is uniquely enabled by direct intratumoral administration as a conduit for lymph node exposure. Current interventional radiology and surgery techniques allow access to almost any surface or visceral lesion, bolstering intratumoral immunotherapies in human cancer as a viable approach. Notably, the FDA-approved oncolytic therapy talimogene laherparapvec (TVEC) is administered intratumorally. In addition, intranodal or s.c. peritumoral administration, which we explore in our αCD45-Cyt paradigm, can be considered as alternative administration routes for TDLN exposure. Finally, while we explored some facets of CD45 biology, there are other potential aspects to consider. Differential splicing of human CD45 can correspond to distinct cell states, with shorter CD45 isoforms upregulated upon CD8 T cell activation(*37*–*39*). A CD45RO specific antibody might therefore enable more targeted delivery of cytokines to activated T cells. In summary, our results highlight CD45 anchoring as a potent modular platform for enhancing the retention and response of immune agonists and provide rationale for further development of CD45 immunocytokines.

## Acknowledgments

We thank the Koch Institute Swanson Biotechnology Center for technical support, specifically the Flow Cytometry and Histology core facilities, as well as the MIT BioMicro center. Figure 1A was partially created with Biorender.

## Funding

NSF Graduate Research Fellowship Program (LS, JRP, WP III, OTP)

Alfred P. Sloan UCEM scholarship (LS)

Ludwig Center Scholarship (LRD)

NCI (P30-CA14051)

NIBIB (EB031082)

Mark Foundation for Cancer Research Aspire Award

Howard Hughes Medical Institute

## Author contributions

Conceptualization: LS, KDW, DJI

Performed Experiments: LS, LM, JRP, JAS, LRD, KQ, WP III, JRS, OTP, RAM

Analyzed Data: LS, YJZ, LM

Writing: LS, KDW, DJI

## Competing interests

LS, KDW, and DJI are inventors on a patent filed by MIT related to the methods in this work.

## Data and materials availability

Materials are available upon request. All sequencing data generated in this study will be made publicly available at time of publication.

## Supplementary Materials

Materials and Methods Figs. S1 to S9

## Materials and Methods

### Cell lines and animals

Cell lines CTLL-2 (ATCC), Expi293F (Gibco), HEK-Blue IL-12 (Invivogen) were cultured according to vendor instructions. MC38 (Kerafast), MC38-ZsGreen (developed in the lab as described previously(*40*)), and B16F10 (ATCC) cells were cultured in Dulbecco’s Modified Eagle’s Medium (DMEM) supplemented with 10% fetal bovine serum (FBS), 100 units ml^−1^ penicillin and 100 μg ml^−1^ streptomycin. All cells were maintained at 37 °C and 5% CO_2_, and all confirmed to be negative for mycoplasma.

All animal studies and procedures were carried out following federal, state and local guidelines under an institutional animal care and use committee-approved animal protocol by the Committee of Animal Care at MIT. Female C57BL/6J (The Jackson Laboratory, 000664) and *Batf3−/−* (The Jackson Laboratory, 013755) mice at 6-8 weeks age were purchased and maintained in the animal facility at MIT.

### Cloning and protein purification

Gene blocks (gBlock, IDT) encoding for the light and heavy chain variable regions of anti-CD45 (clone: 30-F11(*41*)) or untargeted isotype control (anti-FITC, clone: 4-4-20) were cloned into a mouse kappa light chain and IgG2c backbone with LALA-PG(*22*) mutations, respectively, in the mammalian expression vector gWiz (Genlantis). For αCD45-IL15, murine IL-15/IL-15Rα_sushi_ , as described previously(*12*), was then cloned at the C terminus of the anti-CD45 (or anti-FITC control) heavy chain. To generate αCD45-IL12, murine single chain IL-12 (scIL-12), as described previously(*12*), was cloned at the N terminus of the anti-CD45 (or anti-FITC control) heavy chain. For both IL-15 and IL-12 immunocytokinse, a (Gly_4_Ser)_3_ linker was used between the cytokine and antibody. A gBlock encoding for the extracellular domain of mouse CD45RO (used for ELISAs; sequence obtained from Uniprot) was cloned into gWiz with a His_6_ tag. Plasmid sequences confirmed by Sanger sequencing (Quintara Biosciences) were transformed into Stellar Competent Cells (Takara Bio) and purified using the NucleoBond Xtra Midi endotoxin-free midi-prep kit (Takara Bio). For immunocytokines, equal mass of heavy and light chain plasmids were transfected into Expi293F cells (Gibco) per manufacturer’s instructions and harvested 6 days after transfection. All immunocytokines were purified using rProteinA Sepharose Fast Flow resin (Cytiva Life Sciences) and validated for size by SDS-PAGE. His-tagged CD45RO was purified by TALON affinity resin (Takara) according to the manufacturer’s instructions. Purified proteins were confirmed to be endotoxin-free (<0.1 EU per dose) using the Endosafe Nexgen-PTS system (Charles River). Purified proteins were flash-frozen in liquid nitrogen and stored at -80C until use.

### ELISA and bioactivity assays

For ELISA assays, Maxisorp 96-well flat bottom plates (ThermoFisher) were coated with recombinant CD45RO at 0.2µg/mL in PBS (Corning) overnight at 4C. Subsequent washes were performed with PBST (PBS supplemented with 0.05% v/v Tween-20 (Millipore-Sigma)). Blocking was performed in PBSTA (PBS supplemented with 1% w/v BSA (Sigma Aldrich) and 0.05% v/v Tween-20) overnight. Immunocytokines were diluted to relevant concentrations in PBSTA and detected via an HRP-conjugated anti-mouse IgG secondary diluted 1:3000 (Abcam). 1-Step Ultra TMB-ELISA substrate solution (ThermoFisher) was added to develop for 5 minutes and quenched with 2N sulfuric acid (VWR). Absorbance at 450 nm with reference at 570nm was measured on an Infinite M200 microplate reader (Tecan).

For IL-15 bioactivity, 10,000 CTLL-2 cells were seeded in a 96-well U-bottom plate in incomplete media per manufacturer’s instructions with stated IL-15 immunocytokine dilutions for 48hrs at 37C. Proliferation was measured via CellTiter-Glo 2.0 Assay (Promega) following manufacturer’s instructions. Luminescence was measured on a microplate reader (Tecan) with an integration time of 0.25 seconds. IL-12 bioactivity was measured with HEK-Blue IL-12 reporter cells according to manufacturer’s instructions (Invivogen).

### Primary CD8^+^ T cell preparation

Spleens from 6-8 week old C57BL/6J female mice were harvested and processed into single-cell suspensions. CD8^+^ T cells were isolated using the EasySep Mouse CD8^+^ T cell isolation kit (StemCell Technologies) and resuspended at a concentration of 10^6^ cells/mL in complete RPMI supplemented with 1X sodium pyruvate (ThermoFisher), 1X non-essential amino acids (ThermoFisher), and 1X beta-mercaptoethanol (ThermoFisher). Media was additionally supplemented with 10 ng/mL murine IL-2 (BioLegend) prior to resuspension and subsequent passaging. Isolated CD8+ T cells were activated for 48 hours on 6-well non-tissue culture (non-TC) treated plate that was precoated with 0.5 µg/mL anti-CD3 (BioXCell, Clone 2C11) and 5 µg/mL anti-CD28 (BioXCell, Clone 37.51) overnight at 4C. The plate was washed twice prior to activation. Following activation, T cells were cultured for 48 hours before use in downstream experiments.

### Fluorescence-quenching internalization assay

Internalization assays were performed and analyzed as described previously(*42*). Briefly, immunocytokines were conjugated with AlexaFluor 488 (AF488) using NHS ester chemistry (Invitrogen). Free dye was removed by Zeba spin desalting column purification (ThermoFisher). 100,000 primary CD8 T cells were seeded in 96-well plate and incubated with AF488-labeled immunocytokines at 10 µg/mL staggered at desired time points. Wells were then split such that one set was incubated with 25 µg/mL anti-AF488 quenching antibody for 25 minutes. For human CD8^+^ T cell internalization assay, anti-human CD45 (clone: MEM-28) conjugated to AF488 (ThermoFisher) was used. Human CD8+ T cells were purchased from StemCell Technologies. Viability was assessed via DAPI staining. AF488 signal was measured using a BD LSR Fortessa and data were analyzed in Flowjo.

### Analysis of STAT phosphorylation by flow cytometry

For *in vitro* STAT5 experiments, primary CD8^+^ T cells cultured as described above were starved of IL-2 for 24 hours and seeded into 96-well plates at 100k cells/well. IL-15 immunocytokines were added at 10 µg/mL for 1 hour and subsequently washed with incomplete T cell media (no IL-2) twice prior to resting for 24 hours. IgG-IL15 added at the same molar concentration (without washing) was used as a continuous control. Cells were immediately fixed in media with equal volumes of BD Phosflow Fixation Buffer I at 37C for 10 minutes. When required, cells were stained with Zombie Aqua viability stain for 5 minutes in PBS (1:1000 dilution) prior to fixation. Cells were permeabilized for 30 minutes on ice with BD Phosflow Perm Buffer III that had been pre-chilled to -20C. Staining with anti-pSTAT5 antibodies (clone 47, BD) conjugated to AF647 or PE was carried out at room temperature for 1 hour. STAT4 experiments were performed identically, but used complete media supplemented with IL-2 and immunocytokine incubation was performed at 2 µg/mL for 20 minutes. pSTAT4 signal was detected with anti-pSTAT4 (clone 38, BD). For trans-signaling experiments, immunocytokines were conjugated with AF647 using NHS-ester chemistry. Preloaded cells were labeled with CFSE per manufacturer’s instructions.

Measurement of STAT5 levels *in vivo* was carried out as previously described(*43*). Briefly, TDLN were processed into single cell suspensions directly in BD Fixation Buffer I and samples were incubated at 37C for 10 minutes. Downstream permeabilization and staining was performed as described above. In all cases, pSTAT signal was measured using a BD LSR Fortessa and data were analyzed in Flowjo.

### Tumor inoculation and treatment preparation

For all single-tumor experiments, mice aged 6-8 weeks old were injected subcutaneously in the shaved right flank with 1M tumor cells (MC38, MC38-ZsGreen, or B16F10) in a volume of 50 μL PBS. For two-tumor experiments, the contralateral tumor was inoculated on the left flank 3 days after the primary tumor, as stated in study schematics. Prior to treatment, mice were randomized to ensure equal mean initial tumor size across groups. Immunocytokines were prepared at their stated doses (1μg for IL-12 immunocytokines, 10μg for IL-15 immunocytokines, where the mass indicated is the mass of the entire fusion protein) and dosed intratumorally in 20 μL PBS unless otherwise stated. Doses were informed by our biodistribution experiments as well as previous intratumoral cytokine work from our lab(*11*). Peritumoral administration was performed in 50 μL PBS injected s.c. at the tail-base. Intraperitoneal administration was performed in 100 μL PBS. Tumor area was calculated as the product of tumor length and width. For single-tumor studies, mice were euthanized when tumor area exceeded 100 mm^2^; for two-tumor studies, mice were euthanized when cumulative tumor area exceeded 200 mm^2^. Immune memory rechallenge experiments were carried out 100 days after initial challenge with 10^5^ tumor cells on the left flank. Age-matched naïve mice were used as controls for these studies. For the lung metastasis model used in Extended Data Fig. 6, 0.2M B16F10 cells in 100 μL PBS were inoculated retro-orbitally on the same day as the standard 1M tumor cell flank inoculation.

### Tissue processing for flow cytometry

B16F10 or MC38 tumors were harvested, weighed, and subsequently minced using dissection scissors in gentleMACS mouse tumor dissociation buffer (Miltenyi) prepared per manufacturer’s instructions. As noted in the Miltenyi protocol, Enzyme R was reduced to 20% of the stated amount to preserve surface epitope integrity. Minced tumors were processed on a gentleMACS Octo-dissociator with heaters (Miltenyi) using program mTDK_1 for B16F10 and mTDK_2 for MC38. Dissociated tumors were then filtered through a 70-micron strainer and 25 mg tumor was plated for downstream staining. TDLN were harvested, weighed, and subsequently dissociated and filtered through a 5 mL round-bottom tube with cell-strainer cap (Falcon) using the blunt rubber end of a 1mL syringe plunger (Falcon). 5 mg of TDLN was used for downstream staining. Blood was collected by sub-mandibular bleeding into MiniCollect K2-EDTA tubes (Greiner) and red blood cells were lysed using ACK Lysis Buffer (Gibco). When intracellular cytokine staining (ICS) was performed, as in Fig. 5, samples were resuspended and plated in complete RPMI supplemented with 1X sodium pyruvate (ThermoFisher), 1X non-essential amino acids (ThermoFisher), 1X beta-mercaptoethanol (ThermoFisher), and 1X brefeldin A (BioLegend) and allowed to incubate at 37C for 3 hours prior to staining. Precision counting beads (50uL, BioLegend) were added after initial resuspension and used for downstream data analysis. Viability was assessed with Zombie UV or Zombie NIR dyes (BioLegend, 1:1000) in PBS for 20 minutes at room temperature. Subsequent washes and surface staining was performed in PBS supplemented with 1% bovine serum albumin and 2mM EDTA (ThermoFisher). Samples were resuspended in Mouse Fc block Plus (BioLegend) prior to surface staining for 15 minutes. Antibodies against surface targets used as following: CD3 (17A3, BD, 1:100), CD4 (GK1.5, BD, 1:100), CD8 (53-6.7, BD, 1:200), CD19 (6D5, BioLegend, 1:100), CD24 (M1/69, BioLegend, 1:100), CD25 (PC61, BioLegend, 1:100), CD44 (IM7, BD, 1:100), CD45.2 (BD, clone: 104, 1:100), NK1.1 (PK136, BioLegend, 1:100), MHCII (M5/114.15.2, BioLegend, 1:100), Ly6C (HK1.4, BioLegend, 1:100), F4/80 (BM8, BioLegend, 1:100), PD1 (29F.1A12, BioLegend, 1:100), TIM3 (RMT3-23, BioLegend, 1:100). P15E tetramer (MBL) staining was performed in the presence of 50 nM dasatinib at a 1:75 dilution and anti-CD8 antibody clone KT15 (ThermoFisher) was used to minimize background signal. Dasatinib incubation was not included in the staining mixture for the RNAseq experiment. When performing intracellular staining, cells were fixed and permeabilized using the Foxp3 transcription buffer set (eBioscience). Samples against intracellular antigens used as following: TCF1 (C63D9, Cell Signaling Technologies, 1:250), IFNg (XMG1.2, BioLegend, 1:200), Granzyme B (QA16A02, BioLegend, 1:200). Intracellular staining was performed overnight at 4C. Cells were collected using a BD FACSymphony A3 and data were analyzed in FlowJo.

### Antibody-mediated cellular depletion

Immune cell depletions were carried out with antibodies targeting CD8a (BioXcell, Clone 2.43, 400 μg twice weekly), NK1.1 (BioXCell, clone PK136, 400 μg twice weekly), CSF1R (BioXCell, clone AFS98, 300 μg every other day) as previously described(*12*). All depletions were given i.p. in 100 μL of PBS. Depletions were initiated 1 day prior to treatment and carried out for four weeks. Depletions were carried out in C57BL/6J mice unless otherwise noted.

### Biodistribution serum measurements

IL-15 immunocytokines were labeled with AF647 using NHS-ester chemistry per the manufacturer’s instructions. Free dye was removed with Zeba desalting columns. Molar amounts of dye for each immunocytokine were matched prior to dosing. Serum concentrations were calculated based on a standard curve prepared using known amounts of serially diluted AF647-labeled immunocytokine.

### FTY720 Preparation and dosing

FTY720 hydrochloride (Sigma Aldrich) was stored in stock solutions at 10 mg/mL in DMSO. Prior to treatment, stock solutions were diluted to a dose of 30μg in 150 μL in PBS. In 2-tumor MC38 studies, FTY720 was dosed every other day i.p. starting on day 5 after tumor inoculation.

### Immunofluorescence Staining

Inguinal LNs and tumors were harvested 24 h post injection as described in Fig. 2A, embedded in O.C.T. buffer (Fischer Scientific) and fresh frozen. 10 μm tissue sections were then post-fixed with 4% paraformaldehyde for 10 minutes, followed by three washes with PBS. Sections were incubated with Fc receptor blocker (Innovex) for 30 minutes and blocked for 1 hour with 5% goat serum, 2.5% BSA in PBS. Staining with primary antibodies was performed overnight at 4°C in blocking buffer (LN: IgD (Biolegend, 405705), and CD8 (Abcam, ab217344). Tumors: CD8, and F4/80 (Abcam, ab105156)). After 3 washes with PBS, the sections were incubated with fluorochrome-conjugated secondary antibody (ThermoFisher, 35551) in blocking buffer for 30 minutes at room temperature. After three washes with PBS, the sections were mounted onto glass slides using mounting media (ProLong Diamond Antifade Mountant, Thermo Fisher Scientific). High magnification images were acquired using a Leica SP8 laser-scanning confocal microscope equipped with a white light laser, a 405 solid state laser line, and selective emission filters. Images were collected using a 25x water immersion lens and a 63x oil immersion lens.

### Immunohistochemistry staining of lung sections

Animals were euthanized and transcardially perfused with PBS before harvesting the lungs. Tissues were fixed overnight in 4% paraformaldehyde (PFA) at 4°C, processed using conventional methods, embedded in paraffin, and sectioned at 10 μm. Sections were then stained with hematoxylin and eosin and scanned using the Aperio Brightfield (Leica Biosystems) Slide Scanning System. The lung tissue and metastatic lesions were automatically detected via distinct pixel classifiers using QuPath v0.4.3.

### CD8 T cell RNA sequencing, mapping, and analysis

For RNA-seq experiments, 2-40k live CD3^+^CD8^+^ CD44^+^p15E^+^ cells processed from TDLN were sorted using a Sony MA900. RNA extraction was performed using the Qiagen RNEasy Micro kit per manufacturer’s instructions. RNA libraries were prepared using the Clontech SMARTer Stranded Total RNA-Seq Kit - Pico Input Mammalian and sequenced using the Illumina NextSeq500 75nt kit. RNAseq reads were aligned to the mouse genome with STAR (v2.7.9a) using ensembl GRCm39 primary assembly as the reference. Aligned reads were quantified using RSEM (v1.3.1) with ensembl GRCm39 (release 110) transcript annotations. The resulting counts were analyzed in R using DESeq2 for differential expression analysis, fgsea for gene set enrichment analysis (GSEA), and msigDB for the gene set database. Data visualization was done with ggplot2 and ComplexHeatmap. GSEA was utilized for the correlative analysis between our RNAseq data and the gene expression signatures from Prokhnevska et al. 2023. The gene expression count matrix was obtained from NCBI Gene Expression Omnibus with accession No. GSE216731. Differential gene expression analysis was performed on the LCMV Arm and TDLN groups, and the genes were ranked by Wald test statistics. The ranked genes were compared to gene signatures from our data, specifically, the upregulated genes in αCD45-Cyt vs. untreated mice and the upregulated genes in untreated vs αCD45-Cyt mice. The enrichment score suggests the degree of correlation with T cells from either LCMV Arm or TDLN.

### Statistical Methods

Statistics were computed in GraphPhad Prism v9 as denoted in the figure captions. For *in vitro*, biodistribution, and flow cytometry immunophenotyping experiments, comparisons were made by *t*-test or one-or two-way analysis of variance (ANOVA) followed by Tukey’s multiple comparison test. Survival comparisons were made by log-rank (Mantel-Cox) tests. Differential gene expression analysis in the RNA-seq data was performed by Wald tests. In all RNA-seq analysis, P values are corrected by Benjamini-Hochberg to account for multiple hypothesis testing. Exact P values are denoted in figures.

**Fig. S1.**
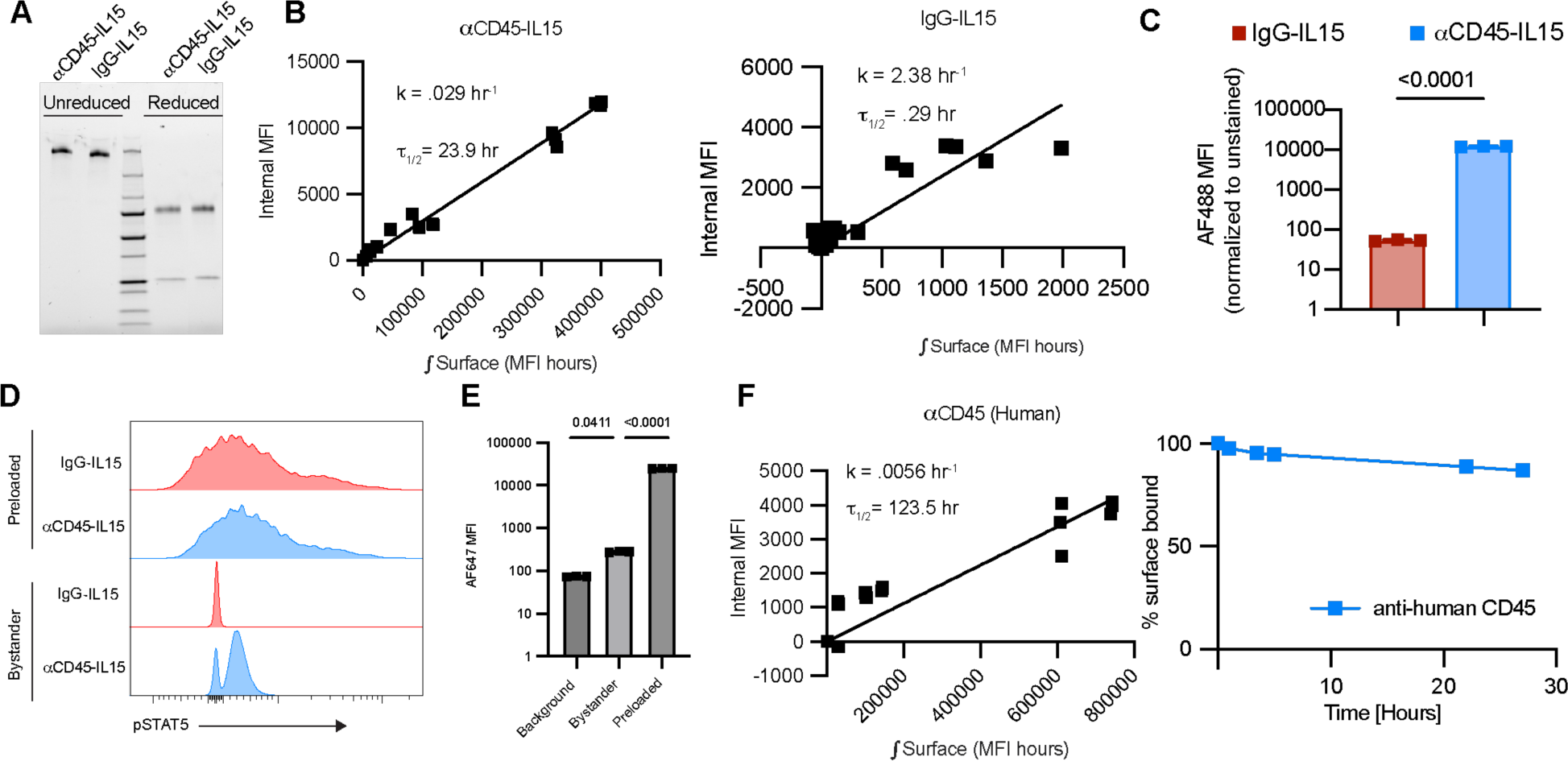
αCD45 immunocytokines show prolonged retention on cell surfaces. **(A)** SDS-PAGE of IL15 immunocytokines. **(B)** Derivation of internalization rate and half-life for αCD45-IL15 (left) and IgG-IL15 (right) on primary CD8 T cells (n = 3 biological replicates per time point). Surface signal was calculated by fluorescence quenching with an αAF488 antibody. **(C)** AF488 signal of CD8^+^ T cells (n = 3 biological replicates) stained for 25 minutes with AF488-labeled αCD45-IL15 or IgG-IL15 for internalization experiment. Unstained background signal was subtracted for analysis. **(D)** Representative pSTAT5 staining (n = 3) for the experimental set-up described in Fig. 1f. **(E)** AF647 MFI signal of cells incubated with αCD45-IL15 (preloaded) and mixed with bystander cells. **(F)** Derivation of internalization rate (left) and internalization kinetics (right) of primary human CD8^+^ T cells incubated with anti-human CD45 labeled with AF488 for stated time points. Surface signal was calculated by fluorescence quenching with an αAF488 antibody. All plots show mean ± SD. P values were determined by t-test or one-way ANOVA followed by Tukey’s multiple comparison test.

**Fig. S2.**
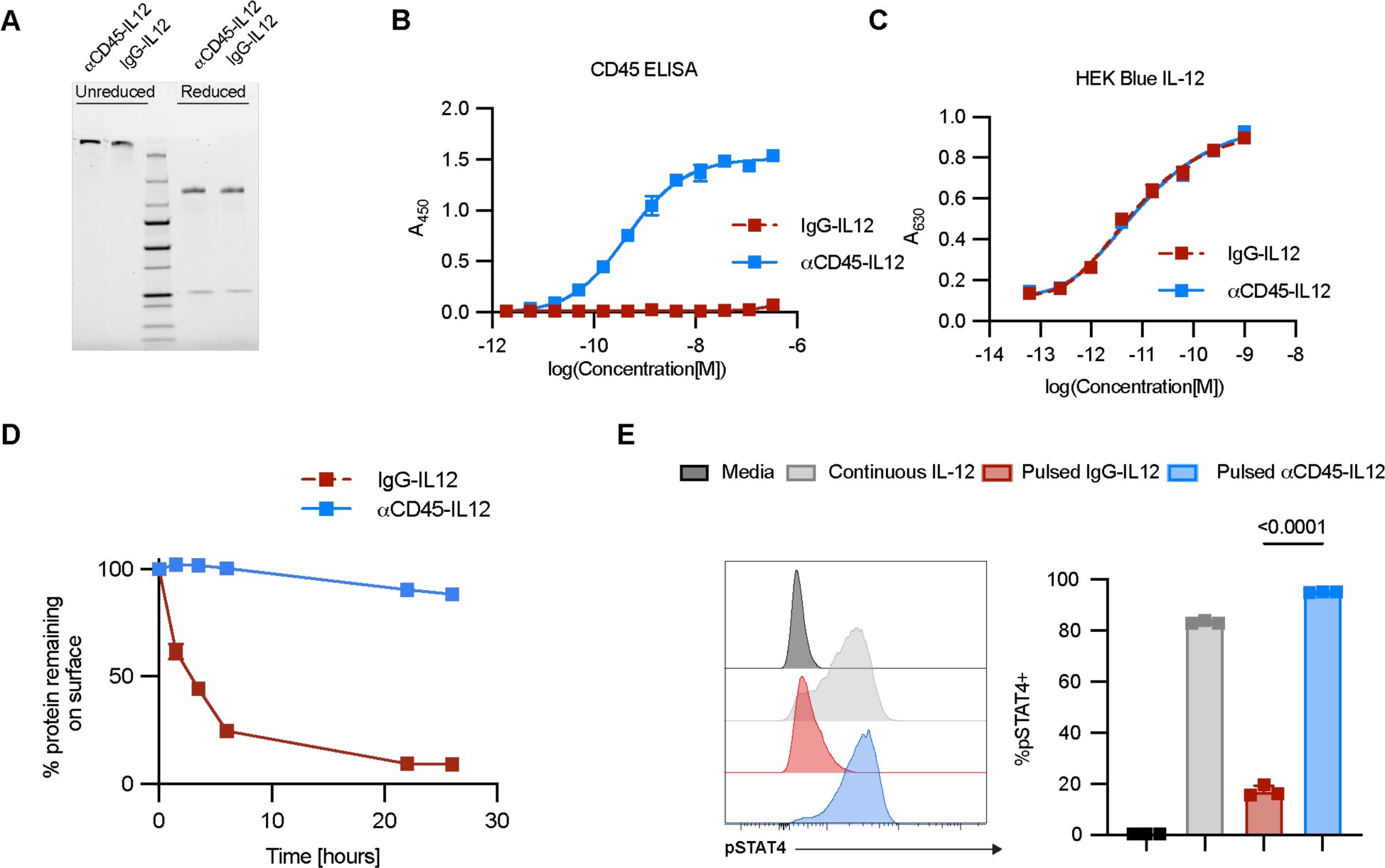
CD45-IL12 immunocytokines are retained on the cell surface and display extended signaling. **(A)** SDS-PAGE of IL12 immunocytokines. **(B)** ELISA absorbance measurement of αCD45-IL12 and IgG-IL12 binding to plate-bound mouse CD45. **(C)** Absorbance measurement of HEK-Blue-IL12 cells following 24 hr incubation with IgG-IL12 or αCD45-IL12 at indicated concentrations. **(D)** Internalization kinetics of IgG-IL12 or αCD45-IL12 labeled with AF488 following binding to primary activated CD8^+^ T cells. Surface signal was calculated by fluorescence quenching with an αAF488 antibody. **(E)** Primary CD8^+^ T cells were pulsed for with IL-12 immunocytokine fusions for 20 minutes. Cells were washed to remove unbound cytokine, and 24 hours later fixed, permeabilized, and stained for flow cytometry analysis of pSTAT4. All plots show mean ± SD from n = 3 biological replicates. P values were determined by one-way ANOVA followed by Tukey’s multiple comparison test.

**Fig. S3.**
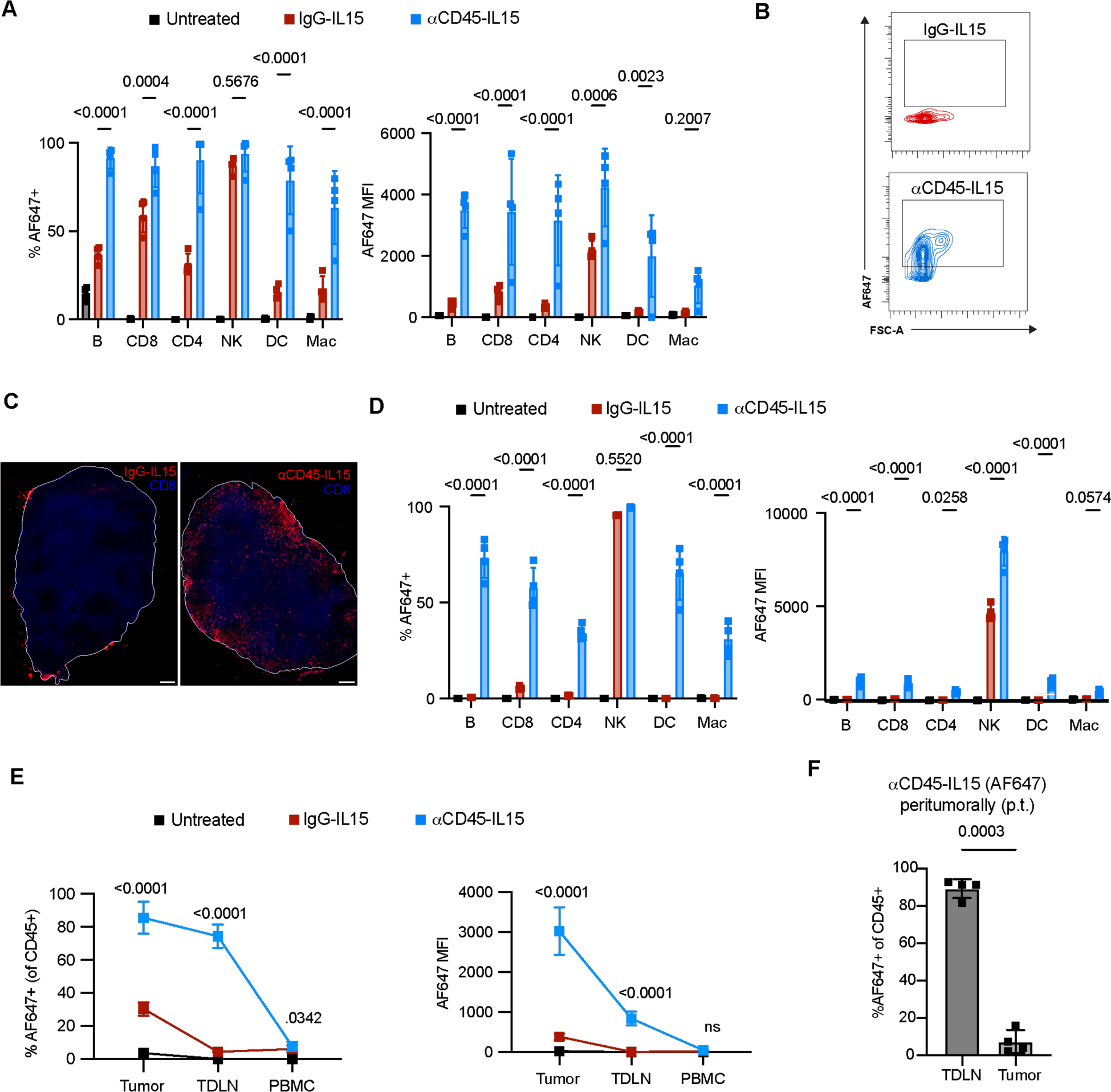
αCD45-IL15 displays altered biodistribution compared to non-targeted IgG-IL15. (**A** to **E**), AF647-labeled αCD45-IL15 or IgG-IL15 was dosed (10µg) intratumorally in established MC38 tumors 24 hours prior to flow cytometry analysis (n = 4 mice/group). Shown are (A) Tumor cell-type specific biodistribution by percentage (left) or MFI (right). (B) Representative contour plot of AF647 signal on TDLN CD45^+^ cells. (C) Representative TDLN images from the experiment described in Fig. 2a. Scale bar, 200 µm. (D) TDLN cell-type specific biodistribution by percentage (left) or MFI (right). (E) Compartment specific biodistribution by MFI (left) or percentage (right). **(F)** AF647-labeled αCD45-IL15 was dosed (10µg) peritumorally (s.c. at the tail-base) 24 hours prior to flow cytometry analysis (n = 4 mice/group). All plots show mean ± SD from n = 4 biological replicates. P values were determined by t-test, one-way, or two-way ANOVA followed by Tukey’s multiple comparison test.

**Fig. S4.**
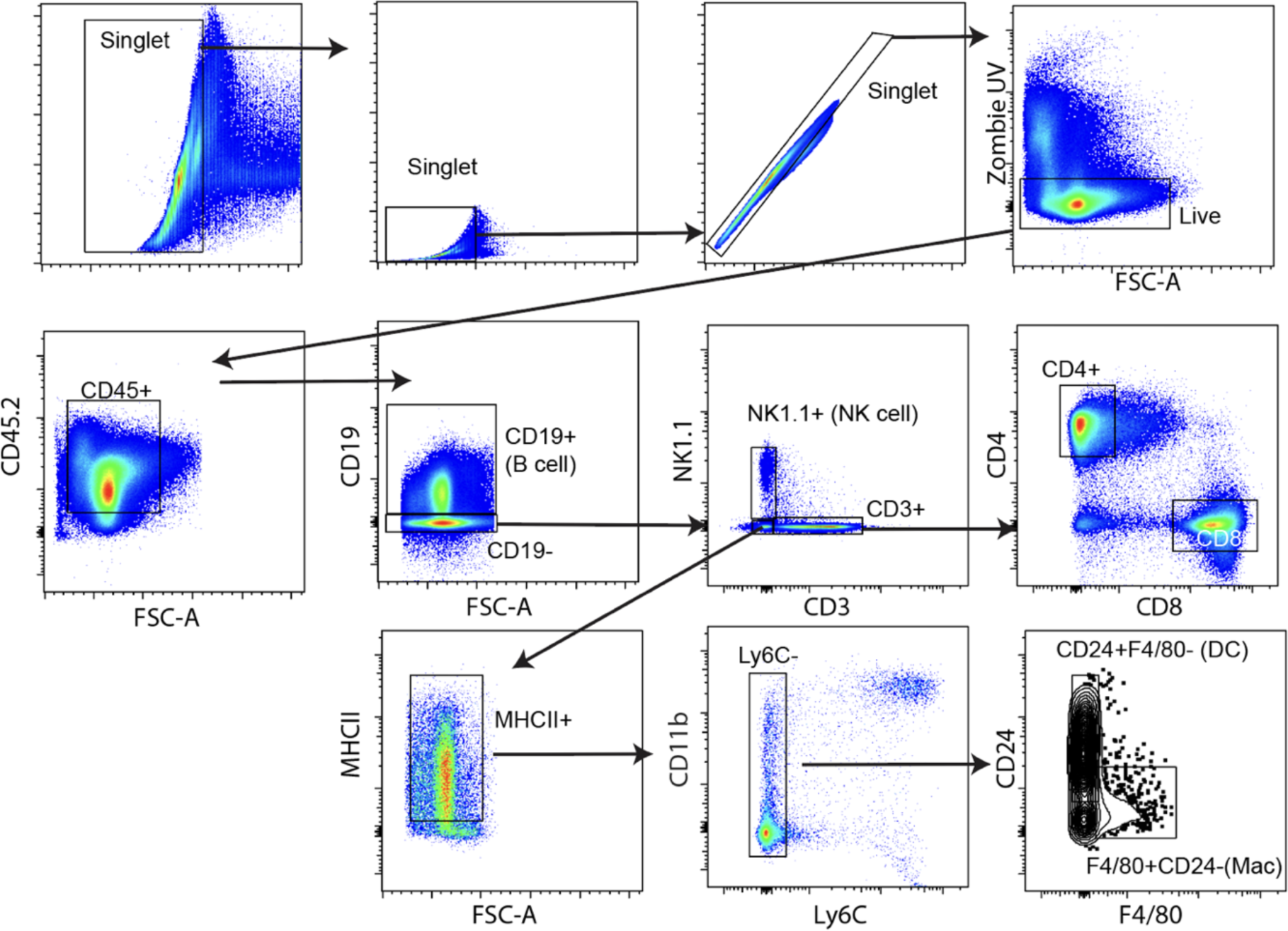
Gating strategy used in Figure 2. Broad leukocyte gating strategy used in Figure 2 and Figure S3.

**Fig. S5.**
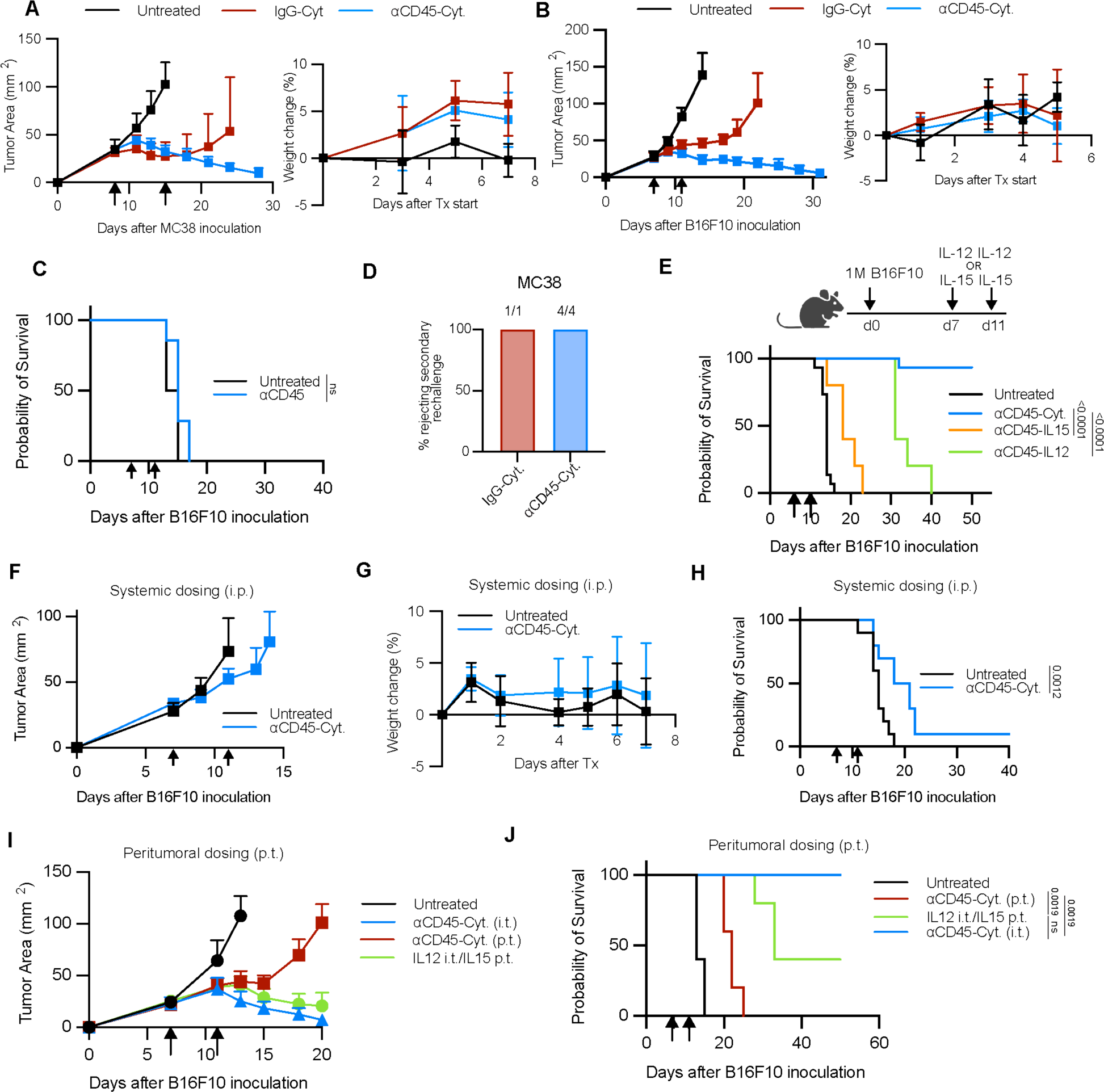
Extended Data Figure 4. αCD45-Cytokine therapy eradicates injected tumors. **(A)** Tumor growth curves (left) and weights (right) of mice (n = 5/group) inoculated with 1M MC38 tumor cells and treated with IgG-Cyt therapy or αCD45-Cyt therapy as described in Fig. 3A. **(B)** Representative tumor growth curves (left) and weights (right) of mice (n = 5/group representative of 3-4 independent experiments) inoculated with 1M B16F10 tumor cells and treated with IgG-Cyt therapy or αCD45-Cyt therapy as described in Fig. 3A. **(C)** Overall survival of mice (n = 6-7/group) bearing B16F10 tumors treated with αCD45 (in the absence of cytokine) on an identical regimen and molar matched dose to that described in Fig. 3A. **(D)** Overall survival of mice cured of MC38 tumors rechallenged with 0.1M MC38 cells 100 days after the initial challenge. **(E)** Kaplan Meier survival plot of mice (n = 5/group) inoculated with 1M B16F10 tumor cells and treated with IL-12 immunocytokine monotherapy (1 µg dose) or IL-15 immunocytokine monotherapy (10 µg dose) as shown on the diagram. **(F)** Tumor growth curves of mice (n = 5/group representative of 2 independent experiments) inoculated with 1M B16F10 cells and treated systemically (i.p.) with αCD45-Cyt therapy with the same doses and schedule as in Fig. 3A. **(G)** Weights of mice shown in (F). **(H)** Overall survival of mice shown in (F). **(I)** Tumor growth curves of mice inoculated with 1M B16F10 cells and treated with the full αCD45-Cyt intratumorally, peritumorally, or the IL-12 immunocytokine given intratumorally and the IL-15 immunocytokine given peritumorally, as described in the text. **(J)** Overall survival of the mice shown in (I). All plots show mean ± SD. P values were determined by log-rank (Mantel-Cox) test.

**Fig. S6.**
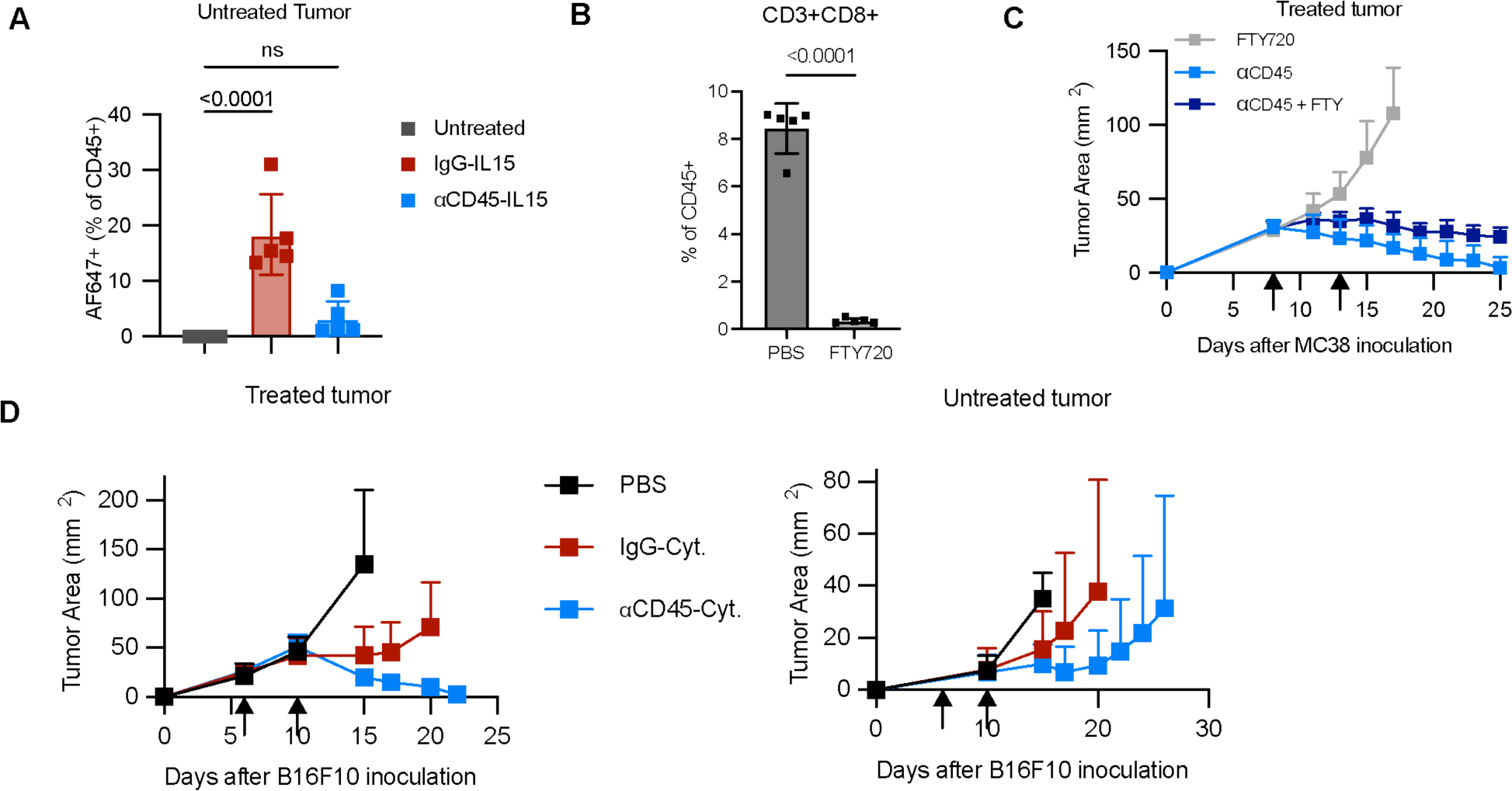
αCD45-Cytokine therapy drives abscopal effects. (A) Mice (n = 5/group) were inoculated with bilateral MC38 tumors as described and dosed with a therapeutic dose of AF647-labeled IL-15 immunocytokine. Shown are AF647 levels on immune cells at the contralateral tumor 24 hours after administration of IL-15. **(B)** Mice (n = 5/group) were dosed i.p. with 1.5mg/kg FTY720. Two days later, PBMCs were collected and shown is the CD8^+^ T cell frequency. **(C)** Mice bearing bilateral MC38 tumors were treated with αCD45-Cyt therapy and 1.5 mg/kg FTY720 i.p. starting on d5 every other day. Shown is the average tumor growth curve for the treated tumor. **(D)** Mice bearing bilateral B16F10 tumors were treated with IgG-Cyt or αCD45-Cyt therapy as described. Shown are tumor growth curves for the treated tumor (left) and contralateral tumor (right). All plots show mean ± SD. P values were determined by t-test or one-way ANOVA followed by Tukey’s multiple comparison test.

**Fig. S7.**
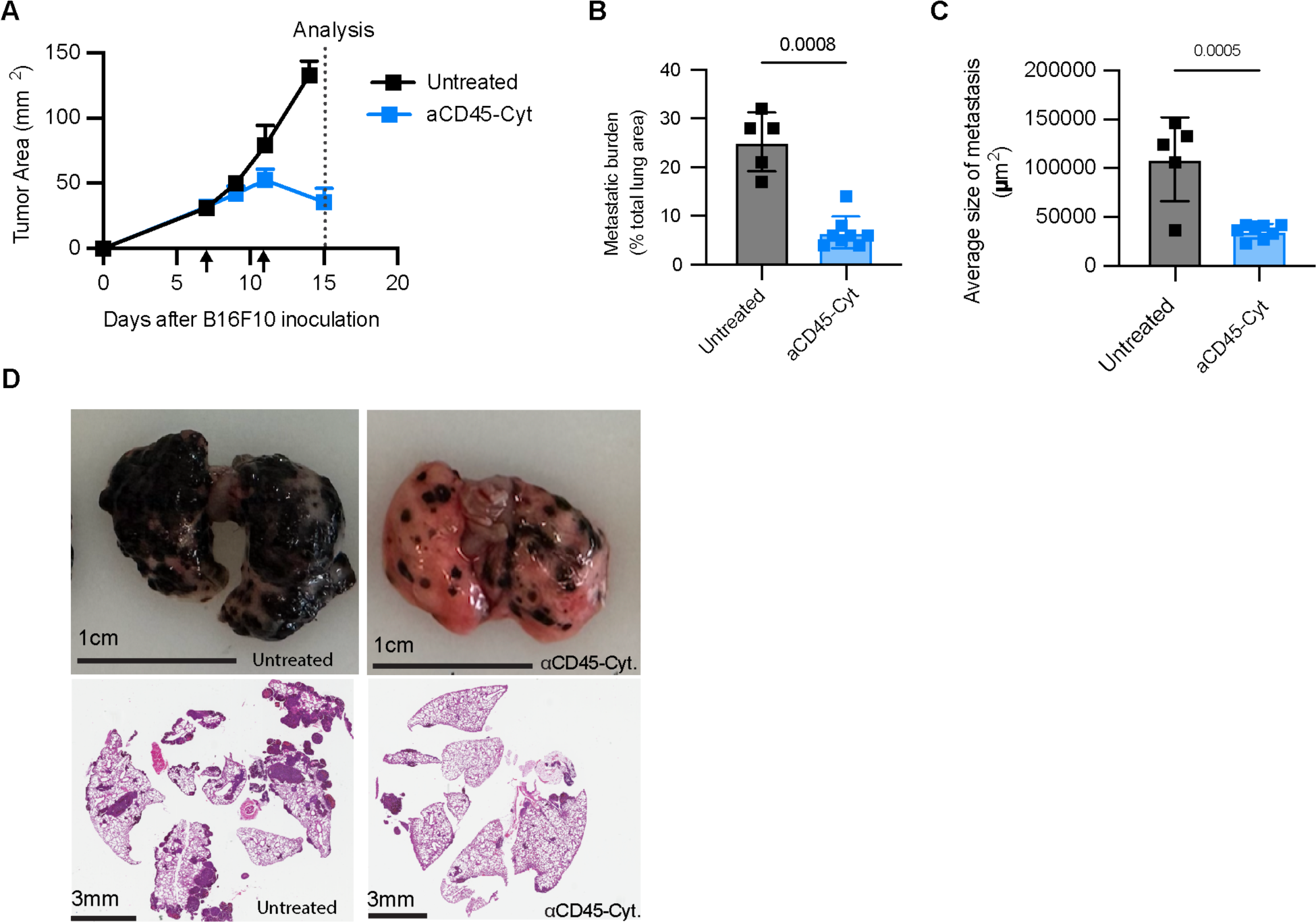
αCD45-Cyt therapy triggers systemic anti-tumor immunity. C57BL/6J (n=5-8/group) were simultaneously inoculated with 1M B16F10 cells s.c. and 0.2M cells retro-orbitally (r.o.) to develop lung metastases. Mice were treated with 1μg αCD45-IL12 immunocytokine on day 7 and 10μg αCD45-IL15 on day 11 (αCD45-Cyt). On d15, animals were euthanized and tumor burden was analyzed. **(A)** Tumor growth curves with arrows denoting treatment. Dashed line denotes analysis timeline. (**B**) Metastatic lung burden shown as percentage of lung tissue after αCD45-Cyt treatment. **(C)** Average metastasis size after αCD45-Cyt treatment. **(D)** Representative images of untreated (top left) or αCD45-Cyt treated (top right) lungs. Representative histology of untreated (bottom left) or αCD45-Cyt treated (bottom right) lungs. All plots show mean ± SD. For metastatic burden, P values were determined by Mann-Whitney non-parametric test. For average metastasis size, P values were determined by *t*-test.

**Fig. S8.**
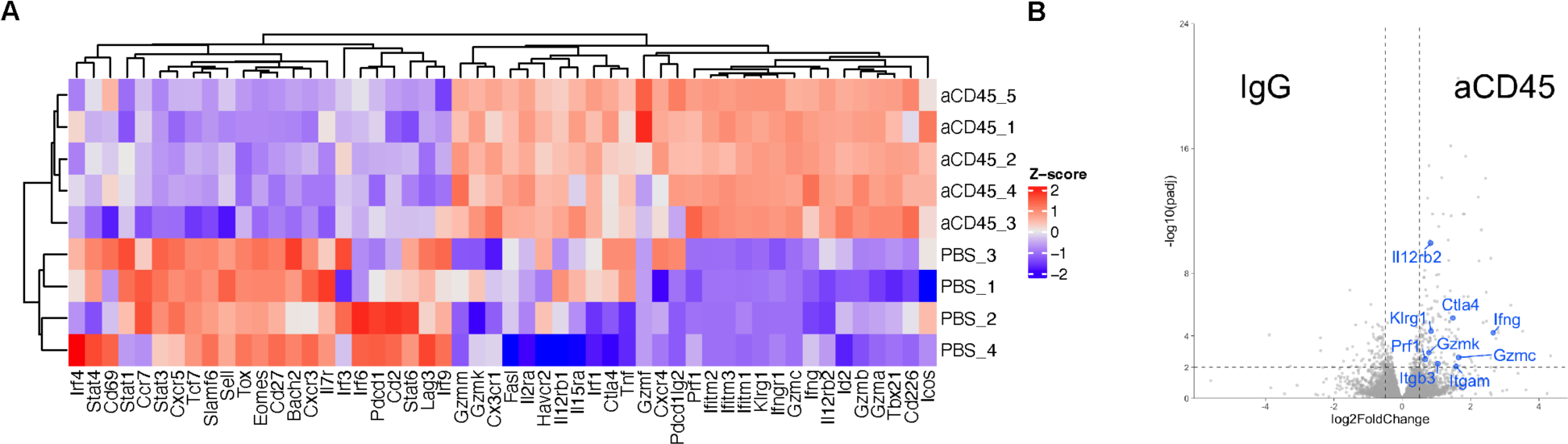
αCD45-Cyt therapy enhances activation of the tumor-specific TDLN compartment. B16F10 tumors (*n* = 4 or 5/ group) treated with αCD45-Cyt or IgG-Cyt therapy were sorted for CD8^+^ T cells specific for the immunodominant p15E retroviral antigen for downstream RNA-seq. **(A)** Differential gene expression of select inflammatory and activation markers between untreated and αCD45-Cyt treated mice. **(B)** Volcano plot of differentially expressed genes between IgG-Cyt and αCD45-Cyt with selected genes labeled.

**Fig. S9.**
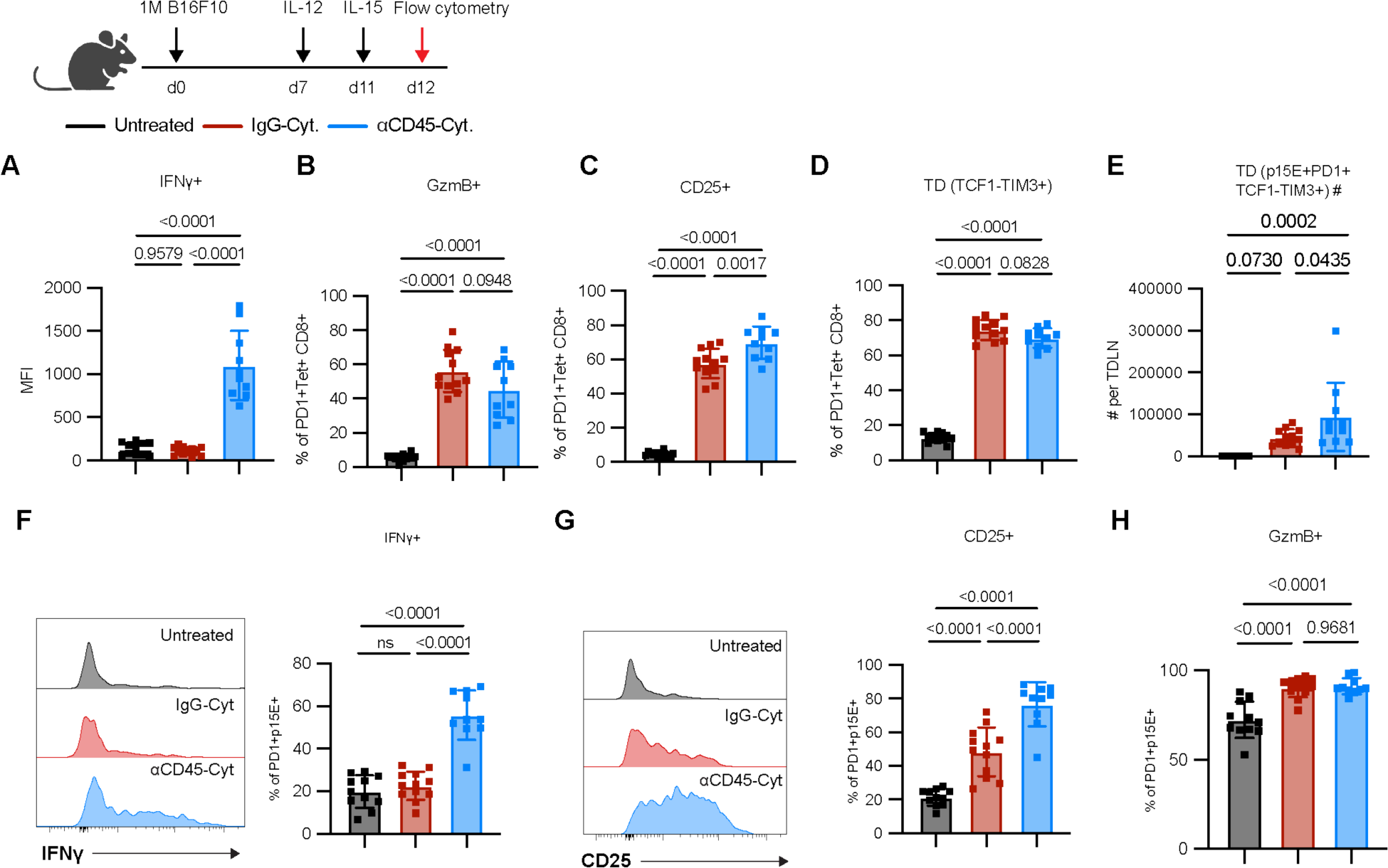
αCD45-Cyt therapy elicits potent effector phenotype on tumor-specific CD8^+^ T cells within the TDLN and TME. Mice (n = 10-12/group across two independent studies) inoculated with 1M B16F10 tumor cells were treated with IgG-Cyt or αCD45-Cyt therapy on d7. Lymph nodes and tumors were harvested 24 hours after completion of therapy. (**A** to **E**) demonstrate the effects of αCD45-Cyt therapy on p15E^+^CD8^+^ T cells in the TDLN. Shown are (A) IFNg MFI levels, (B) Granzyme B expression, (C) CD25 expression, (D) Expression of terminally differentiated phenotype (TCF1^-^TIM3^+^), (E) Absolute TDLN counts of terminally differentiated cells.(**F** to **H**) demonstrate the effects of αCD45-Cyt therapy on intratumoral p15E^+^CD8^+^ T cells. (F) Sample histogram of IFNg staining (left) and expression data (right). (G) Sample histogram of CD25 staining (left) and expression data (right). (H) Granzyme B expression. All plots show mean ± SD. P values were determined by one-way ANOVA followed by Tukey’s multiple comparison test.

